# Biochemical Characterization of Fatty Acid Thioesterase Target Site Mutants and their Implication on Herbicide Resistance

**DOI:** 10.64898/2026.06.11.731613

**Authors:** Paula Wagner, Jens Lerchl, Michael Betzt, Aimone Porri

## Abstract

Herbicide resistance threatens effective weed control in modern agriculture, particularly in grass weeds such as *Alopecurus myosuroides* and *Lolium multiflorum*. Cinmethylin is a pre-emergence herbicide with a novel mode of action that inhibits plastidial fatty acid thioesterases (FATs), enzymes essential for fatty acid biosynthesis. Although no cases of field resistance to cinmethylin have been reported, its resistance risk has not been fully assessed. In this study, we biochemically characterized defined amino acid substitutions in FAT A and FAT B to evaluate their effects on cinmethylin inhibition profile. Some substitutions in FAT A reduced inhibition *in vitro*, with mutations at residue R171 causing the largest shifts in sensitivity. However, these highly resistant variants required multiple specific nucleotide polymorphisms and are therefore predicted to be unlikely to arise in weed populations. In FAT B, sensitivity shifts were generally moderate. Importantly, most substitutions that reduced cinmethylin sensitivity also impaired enzymatic activity, suggesting limited viability *in planta*. Overall, these results indicate that while theoretical target-site resistance mechanisms exist, the practical risk of rapid resistance evolution to cinmethylin is low, supporting its value for integrated grass weed management

## Introduction

Since the domestication of crops, humans have faced the persistent challenge of protecting agricultural systems from diverse biotic and abiotic stressors (Latham, 2013; Tronsmo et al., 2020). The introduction of synthetic herbicides in the twentieth century revolutionized weed control and substantially increased crop productivity (Powles & Yu, 2010). Owing to their efficiency, selectivity and economic advantages over mechanical or manual weed control, herbicides remain indispensable for global food production (Schulte & Theuvsen, 2015). However, continuous herbicide use has imposed strong selection pressure on weed populations, driving the evolution of resistance. Since the first confirmed case of herbicide resistance in 1957, the number of resistant weed species has increased dramatically and poses a major threat to sustainable agriculture. Currently, resistance to 21 of 31 known herbicide modes of action (MoA) has been documented in 273 species across 75 countries (Heap, 2025). Herbicide resistance arises through genetic variation that enables individual plants to survive and reproduce under herbicide selection pressure. Resistance mechanisms are broadly categorized into non-target-site resistance (NTSR), such as enhanced metabolism or altered translocation, and target-site resistance (TSR), which results from modifications of the herbicide target protein (Gaines et al., 2020). TSR is often conferred by single-nucleotide polymorphisms (SNPs) in the herbicide target gene that reduce binding affinity while maintaining enzymatic function (Gaines et al., 2020; Beckie, 2020; Murphy & Tranel, 2019). Such SNPs can confer resistance to a single herbicide or an entire chemical class, depending on the structural similarity of the compounds and their target interactions. Therefore, early biochemical and structural evaluation of herbicide targets is essential for assessing resistance risk and for guiding the development of sustainable modes of action in weed management.

Grass weeds such as *blackgrass* [*Alopecurus myosuroides* Huds.] and *Italian ryegrass* [*Lolium multiflorum* Lam.] are among the most problematic species in cereal production due to their rapid adaptation and widespread resistance to acetyl-CoA carboxylase (ACCase) and very-long-chain fatty-acid (VLCFA) synthesis inhibitors and ALS herbicides (Messelhäuser et al., 2021; Scarabel et al., 2020). This problem has increased the demand for herbicides with novel biochemical targets and unique MoAs.

Cinmethylin [(1S,2R,4R)-1-methyl-2-[(2-methylphenyl)methoxy]-4-propan-2-yl-7-oxabicyclo[2.2.1]heptane)] represents such an innovation. Discovered by Shell in the 1980s, cinmethylin inhibits fatty acid thioesterases (FATs), key enzymes responsible for terminating plastidial fatty-acid biosynthesis (Campe et al., 2018; Jones et al., 1995). FATs catalyze the hydrolysis of acyl-ACP intermediates and thereby releasing free fatty acids which act as essential precursors for membrane lipids, cutin and suberin (Haslam & Kunst, 2013; Ohlrogge & Browse, 1995). Inhibition of FATs disrupts fatty acid release, leading to impaired lipid metabolism, loss of membrane integrity and ultimately plant death (Campe et al., 2018).

Plant FATs are classified into two families, FAT A and FAT B, which differ in substrate specificity and physiological function and occur in multiple isoforms. FAT A preferentially hydrolyzes unsaturated acyl-ACP substrates, whereas FAT B primarily targets saturated acyl-ACP species (Liu et al., 2022).

Despite its early discovery, cinmethylin was not widely adopted in agricultural practice for several decades due to the dominance of competing herbicide classes targeting ACCase and VLCFA biosynthesis (Campe et al. 2018, Yu et al., 2024). However, the extensive use of these alternative MoAs has resulted in widespread resistance, particularly in problematic grass weeds such as *A. myosuroides.* As resistance to established herbicides increased, the agronomic relevance of cinmethylin was rediscovered (Campe et al., 2018).

For many years, the precise MoA of cinmethylin remained unclear and it was therefore classified by the Herbicide Resistance Action Committee (HRAC) as having an unknown MoA. Recent studies employing cellular target profiling identified FAT isoforms as direct molecular targets of cinmethylin, a finding subsequently confirmed by co-crystallization and biochemical assays using recombinant FAT proteins (Campe et al., 2018). These insights led to the reclassification of cinmethylin as the first herbicide in HRAC Group 30, representing a truly novel MoA. However, the resistance risk associated with this novel target, specifically whether amino acid substitutions in FATs can reduce cinmethylin sensitivity while preserving enzymatic activity, remains insufficiently characterized.

Given the ecological and economic impact of herbicide-resistant weeds it is a key challenge to identify target-site mutations that may confer herbicide resistance. *A. myosuroides* and *L. multiflorum* are among the most problematic grass weeds in Europe (Messelhäuser et al., 2021; Scarabel et al., 2020). The objective of this study was to evaluate the resistance potential of cinmethylin by determining whether defined amino acid substitutions in FATs reduce cinmethylin inhibition while maintaining enzymatic activity. To also assess potential cross-species relevance, FAT sequences from both *A. myosuroides* and *L. multiflorum* were compared to identify conserved regions and candidate positions for amino acid substitutions.

## Materials and Methods

### Study Design

The study was conducted as an exploratory biochemical screening of FAT A and B mutant variants using the *A. myosuroides* backbone sequence. Due to the large number of variants, each mutant was produced once and evaluated under standardized assay conditions alongside wild type controls.

### Expression of recombinant FAT proteins

Putative target-site mutations in FATA and FATB were identified by performing molecular docking simulations in which cinmethylin was computationally positioned within the active binding pockets of both enzymes, allowing the prediction of amino acid residues likely to interact with the herbicide and potentially contribute to altered binding.

WT proteins were expressed in five independent biological replicates; each was assayed in duplicate wells (technical replicate). Mutants were expressed once (one biological replicate) and assayed in duplicate wells.

Recombinant proteins were heterologously expressed in *Escherichia coli (E. coli)..* FAT encoding plasmids (pET24d, c-terminal His-tag) were obtained as synthetic constructs (GeneArt, Thermo Fisher Scientific). Plasmids were transformed into *E. coli* BL21-CodonPlus(DE3)-RIL competent cells (Agilent). Transformants were selected on LB agar containing kanamycin (50 µg mL^-1^) and chloramphenicol (34 µg mL^-1^). Single colonies were used to inoculate LB pre-cultures (20 mL) and incubated at 37°C 140 rpm for 17 h. Main cultures were established in autoinduction medium with ZY-base supplemented with trace elements and carbon sources as described in Tab. 1. Cultures were incubated at 37 °C and 140 rpm for 5 h followed by 25 °C at 140 rpm for 21 h. Cells were harvested by centrifugation (7000 x g, 30 min, 4 °C), pellets were frozen on dry ice and stored at -80 °C until purification.

**Tab. 1.** Media formulations used for recombinant expression of *A. myosuroides* FAT proteins in *E.coli*.

| Component | Final concentration |
| --- | --- |
| LB pre-culture medium |  |
| Tryptone | 10g L<sup>-1</sup> |
| Yeast extract | 5g L<sup>-1</sup> |
| NaCl | 10g L<sup>-1</sup> |
| Kanamycin | 50µg mL<sup>-1</sup> |
| Chloramphenicol | 34µg mL<sup>-1</sup> |
| Autoinduction medium ZY based |  |
| Tryptone | 10.8 g L<sup>-1</sup> |
| Yeast extract | 5.4 g L<sup>-1</sup> |
| MgSO<sub>4</sub> (1 M) | 1 mM |
| Trace elements, including: |  |
| FeCl<sub>3</sub> x 6H<sub>2</sub>O (0,05 M) | 50 µM L<sup>-1</sup> |
| CaCl<sub>2</sub> x 2H<sub>2</sub>O (0,02 M) | 20 µM L<sup>-1</sup> |
| MnCl<sub>2</sub> x 4H<sub>2</sub>O (0,01 M) | 10 µM L<sup>-1</sup> |
| ZnSO<sub>4</sub> x 7H<sub>2</sub>O (0,01 M) | 10 µM L<sup>-1</sup> |
| CoCl<sub>2</sub> x 6H<sub>2</sub>O (0,002 M) | 2 µM L<sup>-1</sup> |
| CuCl<sub>2</sub> x 2H<sub>2</sub>O (0,002 M) | 2 µM L<sup>-1</sup> |
| NiCl<sub>2</sub> x 6H<sub>2</sub>O (0,002 M) | 2 µM L<sup>-1</sup> |
| Na<sub>2</sub>MoO<sub>4</sub> x 2H<sub>2</sub>O (0,002 M) | 2 µM L<sup>-1</sup> |
| 50x5052, containing |  |
| Glycerol | 0.5% (v/v) |
| Glucose | 0.05% (v/v) |
| Lactose | 0.2% (v/v) |
| 50xM, containing: |  |
| NH<sub>4</sub>Cl (2,5 M) | 50 mM |
| KH<sub>2</sub>PO<sub>4</sub> (1,25 M) | 25 mM |
| Na<sub>2</sub>HPO<sub>4</sub> x 2H<sub>2</sub>O (1,25 M) | 25 mM |
| Na<sub>2</sub>SO<sub>4</sub> | 5 mM |
| Kanamycin | 50µg mL<sup>-1</sup> |
| Chloramphenicol | 34µg mL<sup>-1</sup> |
| Pre-culture | 2% (v/v) |

### Protein purification (IMAC) and dialysis

Cell pellets were resuspended in lysis buffer (50 mM KH_2_PO_4_, 150 mM NaCl, pH 7.2) supplemented with BugBuster (10% [v/v]; MilliporeSigma), lysozyme (133 µg mL^-1^), DNAse I (1 mg mL^-1^) and EDTA-free protease inhibitor (cOmplete^TM^, Roche). Lysis was performed for 20 min at room temperature (at 180 rpm). Soluble fractions were purified using Ni-NTA agarose resin (HisPur^TM^ Superflow, Thermo Fisher Scientific) in centrifuge columns (Pierce^TM^, Thermo Fisher Scientific). Columns were equilibrated with lysis buffer, washed with washing buffer (50 mM KH_2_PO_4_, 150 mM NaCl, 20 mM Imidazole, pH 7.2) and proteins eluted with elution buffer (50 mM KH_2_PO_4_, 150 mM NaCl, 200 mM Imidazole, pH 7.2). To remove imidazole, eluates were dialyzed overnight at 4 °C against dialysis buffer (5 mM KH_2_PO_4_, 15 mM NaCl, pH 7.2). Protein concentrations were determined via spectrophotometer (ScanDrop^2^, Analytic Jena). 8-10% (v/v) Glycerol was added prior to storage and samples were stored at -80 °C.

### Fluorescence-based acyl-CoA assay and cinmethylin dose-response

FAT activity was quantified using a fluorescence assay based on release of CoA-SH from acyl-CoA substrates, followed by reaction with the thiol-reactive fluorophore CPM (7-diethylamino-3-(4-maleimidophenyl)-4-methylcoumarin). Reactions were conducted in 96-well plates and measured as endpoint fluorescence (excitation 390 nm, emission 470 nm) using a plate reader (CLARIOstar3 plus, BMG Labtech).

Reactions were conducted in assay buffer (50 mM Tris, 500 mM NaCl, 5 mM EDTA, pH 7.2) Oleoyl-CoA was used as substrate for FAT A and palmitoyl-CoA for FAT B at a final concentration of 100 µM. Cinmethylin stock solutions were prepared in DMSO and serially diluted (1:4) to generate final test concentrations ranging from 1 x 10^-4^ M to 3.81 x 10^-10^ M. Reactions were assembled by combining enzyme, compound (or negative control), substrate and CPM detection reagent with standardized incubation times (compound pre-incubation [600 s], CPM incubation [300 s]). Assay assembly was performed using an automated workstation (Biomek NX^P^,Beckman Coulter) at 10 °C for enzyme and substrate plates. The final enzyme concentration in the assay was 800 nM. Each variant was measured in two technical replicate wells.

### SDS-PAGE verification

Selected purified proteins were analyzed by SDS-PAGE using 10% gels (Mini-PROTEAN TGX Stain-Free^TM^, Bio-Rad). Samples were denaturated (95 °C for 10 min) in LDS sample buffer (NuPAGE^TM^, Invitrogen) and run at 139 V for 60-90 min in Mini-PROTEAN Tetra system (Bio-Rad). Gels were imaged with ChemiDoc^TM^ MP system (Bio-Rad).

### Data processing and statistical analysis

Raw fluorescence values were background corrected by subtracting the negative control signal. Inhibition at the highest cinmethylin concentration (10^-4^ M) was calculated as:

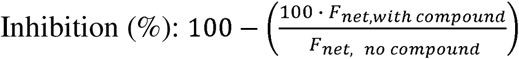

Residual activity (%) was determined relative to WT controls within the same batch:

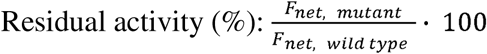

Variants exhibiting <50% residual activity relative to WT were classified as low activity variants and excluded from further sensitivity classification, as such activity levels are assumed to be unlikely to be compatible with plant viability.

Dose-response curves and IC_50_ values were estimated by nonlinear regression using GraphPadPrism (version 10.6.1). IC_50_ values function as comparative screening.

Resistant index was calculated as:

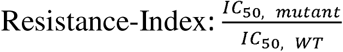

Mutants with IC_50_ values close to WT or below were classified as potentially sensitive to cinmethylin, whereas mutants with increased IC_50_ relative to WT were classified as potentially resistant.

Cross-species sequence analysis (*A. myosuroides* vs. *L. multiflorum*)

Coding sequences of FAT gene from *A. myosuroides* and *L. multiflorum* were aligned to assess sequence conservation and to map potential mutation positions across species. Alignments were performed with Geneious Prime (Biomatters Ltd.).

The mutation probability calculation provides a theoretical estimate of mutation probability based on single nucleotide substitutions and does not account for transition-transversion biases or species specific mutation spectra. Each nucleotide was assumed to mutate with the equal probability into one of the three alternative bases, resulting in a assumed per-base mutation probability of 1/3. At the codon level, three nucleotide positions are available, each with three possible single-nucleotide substitutions which yields a total of nine possible point mutations per codon. If a single point mutation leads to the target codon, the probability was calculated as 1/9; if two distinct point mutations result in the target codon, the probability was calculated as 2/9 and so on.

## Results and Discussion

### FAT A & B WT assay and reference sensitivity

Assay performance and reference sensitivity to cinmethylin were established by using wild type (WT) FAT A and B proteins (n = 5 biological replicates each). FAT A WT showed a mean activity of 31218 ± 2813.3 relative fluorescence units (RFU) and a mean inhibition of 90.5% ± 2.19 at saturating cinmethylin concentration (10^-4^M) (Fig. 1 and Tab. 3). SDS-PAGE confirmed the expected FAT A band at approximately 39 kDa (Fig. 2). Dose-response curves were generated, and IC_50_-values were estimated with a mean of 3.02 x 10^-8^ M ± 3.88 x 10^-9^ (Fig. 3; Tab. 2).

**Fig. 1.**
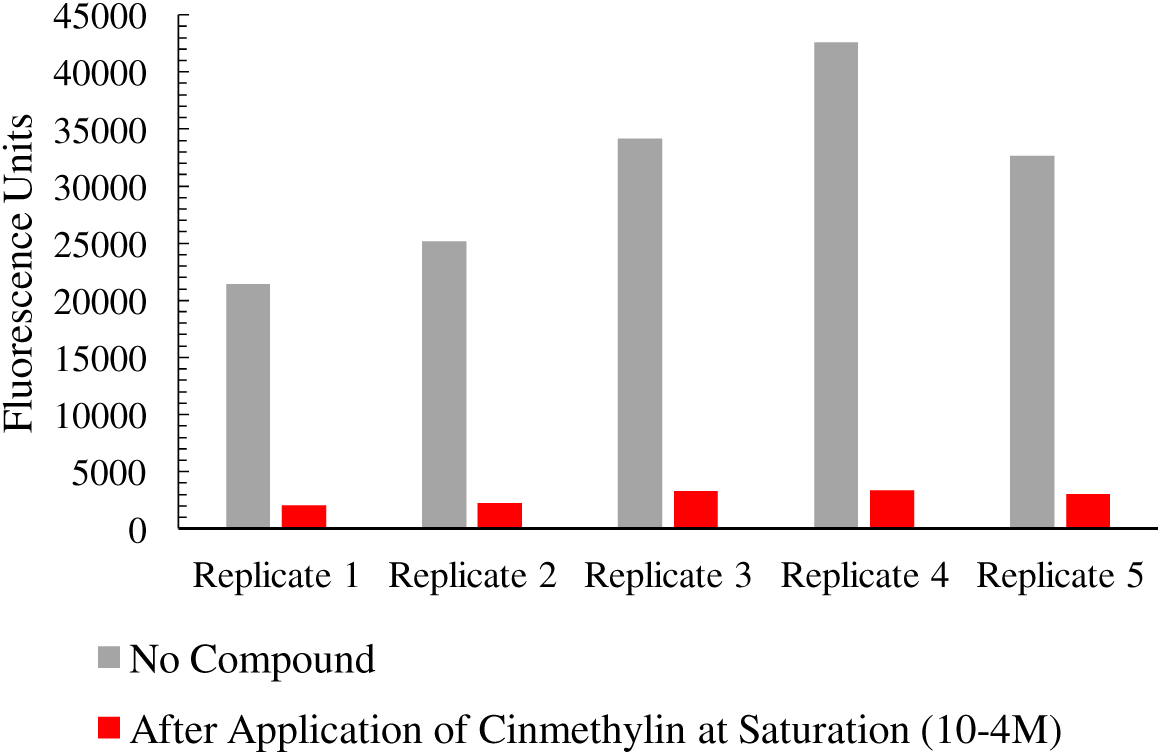
Acyl-CoA Assay of FAT A from *Alopecurus myosuroides* WT biological replicates with and without cinmethylin treatment. Grey bars represent relative in the absence of compound, while red bars show inhibited after application of cinmethylin (10^-4^ M).

**Fig. 2.**
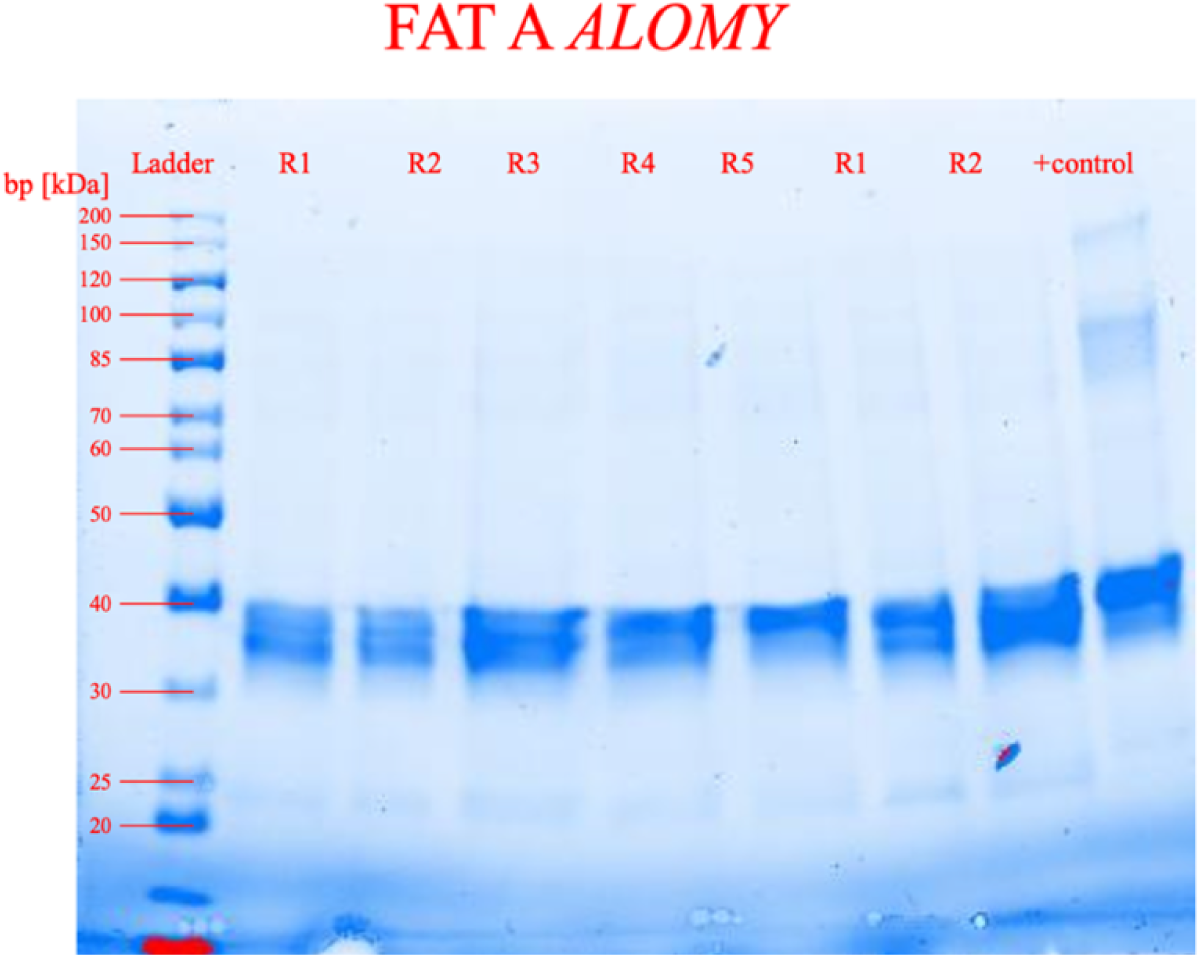
SDS-PAGE of *Alopecurus myosuroides* FAT. **A** with five replicates (including two loaded in duplicate). As a positive control an independently and elaborately purified FAT A protein.

**Fig. 3.**
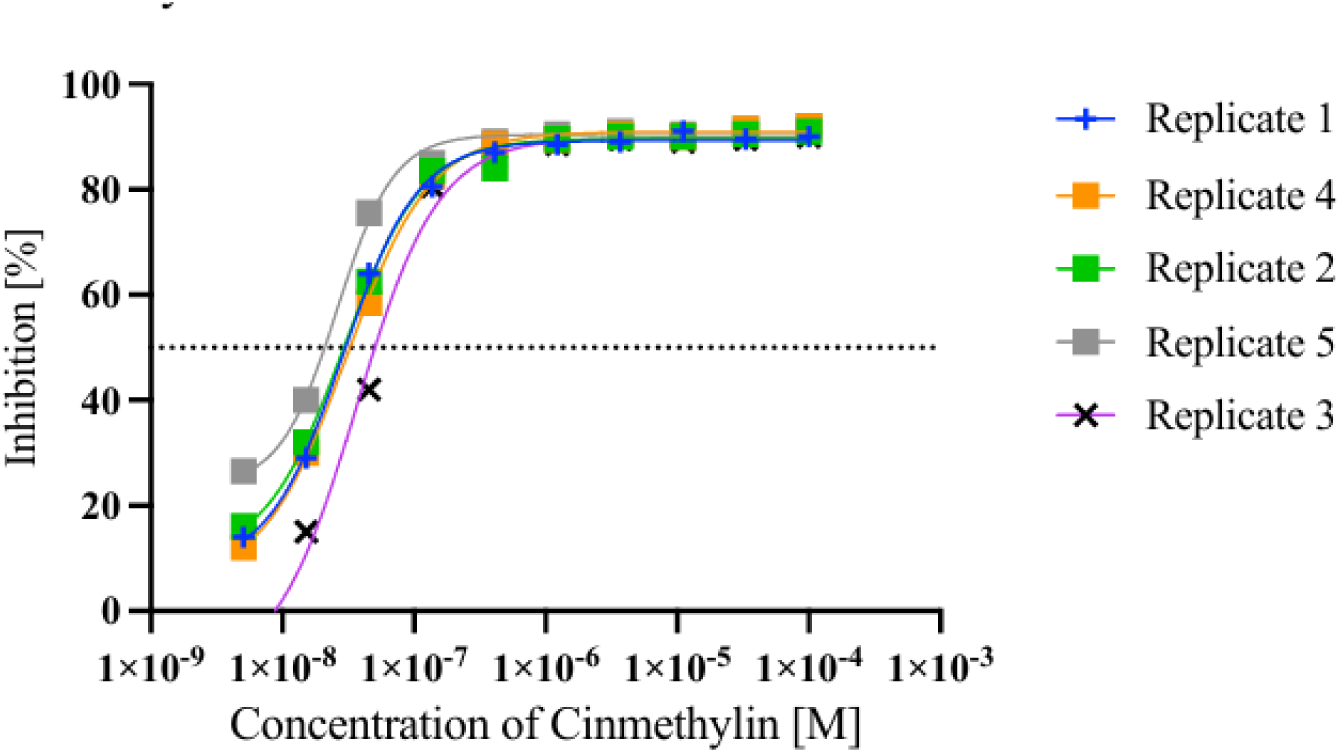
**Determination of IC50 values for *Alopecurus myosuroides* FAT A WT in response to cinmethylin**

**Tab. 2.** IC50 values of *Alopecurus myosuroides* FAT A and B wild-type proteins determined from five biological replicates each.

| Replicate | IC<sub>50</sub> [M] |
| --- | --- |
| FAT A WT 1 | 2,89 x 10<sup>-8</sup> |
| FAT A WT 2 | 2,91 x 10<sup>-8</sup> |
| FAT A WT 3 | 3,64 x 10<sup>-8</sup> |
| FAT A WT 4 | 3,07 x 10<sup>-8</sup> |
| FAT A WT 5 | 2,59 x 10<sup>-8</sup> |
| Mean & SD | 3,02 x 10<sup>-8</sup> ± 3,88 x 10<sup>-9</sup> |
| FAT B WT 1 | 1,17 x 10<sup>-7</sup> |
| FAT B WT 2 | 1,06x 10<sup>-7</sup> |
| FAT B WT 3 | 3,10 x 10<sup>-7</sup> |
| FAT B WT 4 | 1,28 x 10<sup>-7</sup> |
| FAT B WT 5 | 1,58 x 10<sup>-7</sup> |
| Mean & SD | 1,64 x 10<sup>-7</sup> ± 0,84 x 10<sup>-7</sup> |

**Tab. 3.** *Alopecurus myosuroides* FAT A and B mutants with residual activity >50 % and corresponding inhibition values at cinmethylin saturation 10^-4^ M.

| Mutant | Residual activity [%] | Inhibition [%] | IC<sub>50</sub> | Resistance-Index |
| --- | --- | --- | --- | --- |
| FAT A |  |  |  |  |
| WT | n.d. | 90.50 | $3,02 \times 10^{-8}$ | n.d. |
| H112Q | 185.02 | 87.87 | $1.28 \times 10^{-7}$ | 4.24 |
| V116L | 50.58 | 81.87 | $4.85 \times 10^{-8}$ | 1.61 |
| F118Y | 236.41 | 96.61 | $6.17 \times 10^{-8}$ | 2.04 |
| F118C | 222.73 | 52.88 | $8.87 \times 10^{-8}$ | 2.94 |
| A124G | 78.32 | 90.13 | $3.44 \times 10^{-8}$ | 1.14 |
| A124S | 92.77 | 91.60 | $3.77 \times 10^{-8}$ | 1.25 |
| T127K | 65.73 | 86.00 | $6.02 \times 10^{-8}$ | 1.99 |
| R171K | 60.87 | 56.56 | $1.56 \times 10^{-5}$ | 516.56 |
| R171Q | 54.72 | 24.32 | n.d. | n.d. |
| R171I | 84.70 | 15.53 | n.d. | n.d. |
| R171M | 57.39 | 10.96 | n.d. | n.d. |
| W173L | 329.68 | 91.63 | $2.05 \times 10^{-7}$ | 6.77 |
| W173G | 50.96 | 79.33 | $3.79 \times 10^{-8}$ | 1.25 |
| FAT B |  |  |  |  |
| WT | n.d. | 91.5 | $1.64 \times 10^{-7}$ | n.d. |
| A181V | 66.07 | 85.96 | $3.17 \times 10^{-7}$ | 1.93 |
| V226C | 125.77 | 87.61 | $9.47 \times 10^{-8}$ | 0.58 |
| A181V&V226C | 79.39 | 89.56 | $1.23 \times 10^{-7}$ | 0.75 |
| V178A | 114.36 | 92.89 | $1.61 \times 10^{-7}$ | 0.98 |
| V178G | 192.44 | 94.90 | $6.24 \times 10^{-7}$ | 3.80 |
| A190E | 146.85 | 93.35 | $4.15 \times 10^{-7}$ | 2.53 |
| P192R | 111.02 | 92.36 | $1.60 \times 10^{-6}$ | 9.76 |

FAT B WT showed a mean activity of 129284.5 ± 9373.3 RFU and a mean inhibition of 91.5 % ± 6.95 at cinmethylin saturation (Fig. 4 and Tab. 3). SDS-PAGE confirmed the expected FAT B band at approximately 47 kDa (Fig. 5). The mean FAT B WT IC_50_ was and of 1.64 x 10^-7^ M ± 0.84 x10^-7^ (Fig. 6; Tab. 2). Together, these results confirm robust assay performance and provide baseline sensitivity values for subsequent mutant screening.

**Fig. 4.**
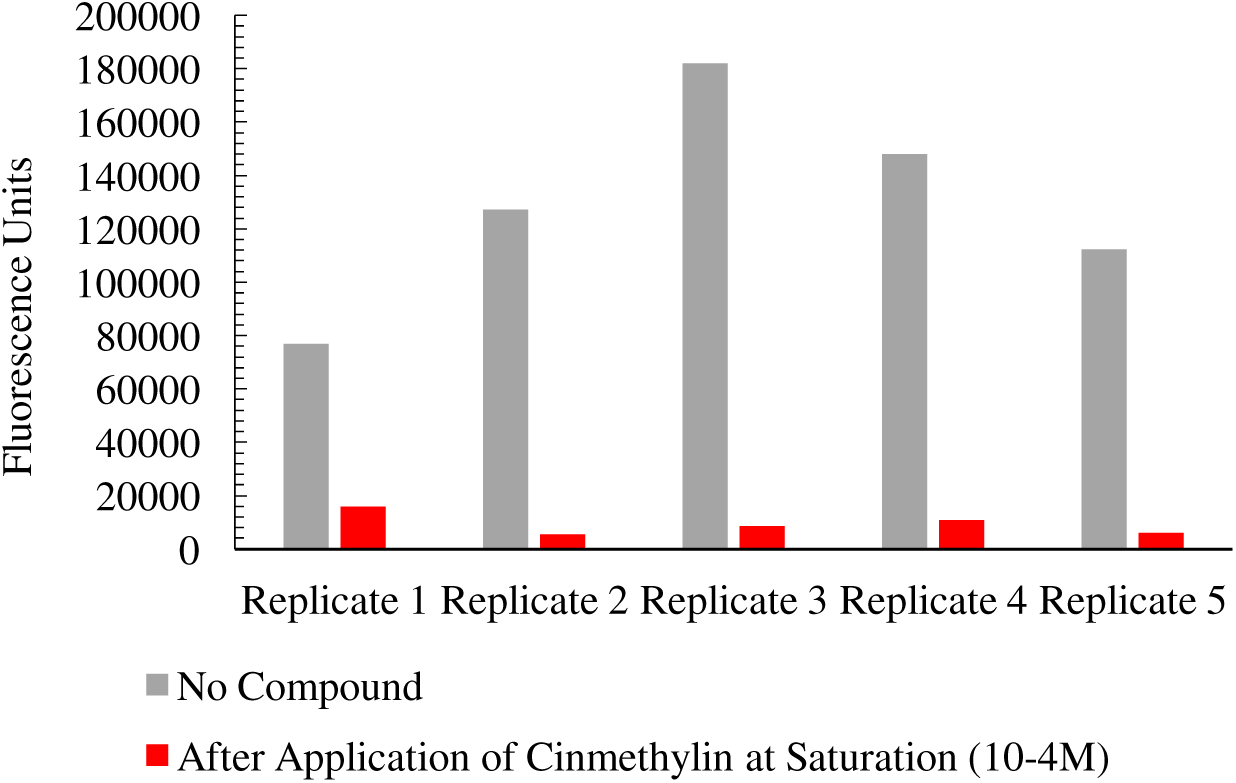
Acyl-CoA Assay of FAT B from *Alopecurus myosuroides* WT biological replicates with and without cinmethylin treatment. Grey bars represent relative in the absence of compound, while red bars show inhibited after application of cinmethylin (10^-4^ M).

**Fig. 5.**
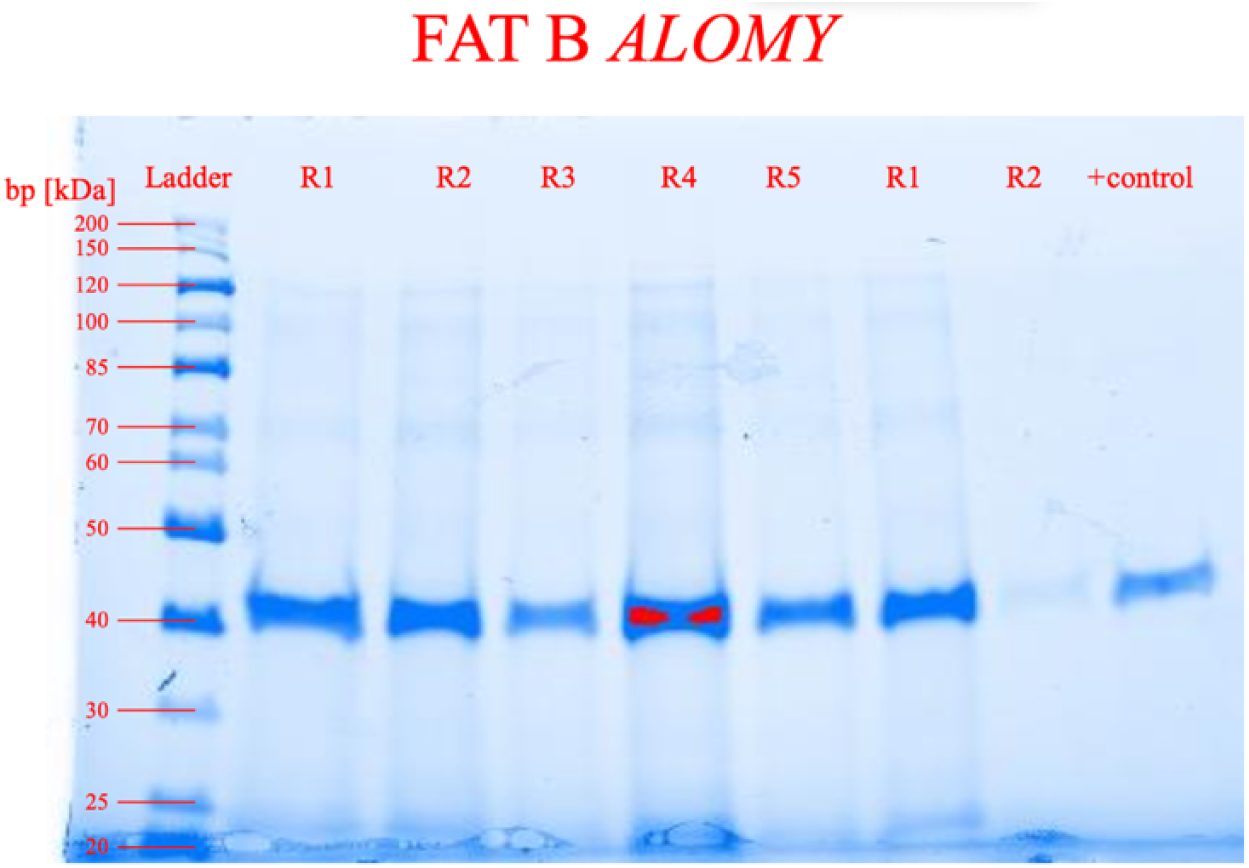
SDS-PAGE of *Alopecurus myosuroides* FAT. **B** with five replicates (including two loaded in duplicate). As positive control an independently and elaborately purified FAT A protein (provided by Marnet, BASF) was included.

**Fig. 6.**
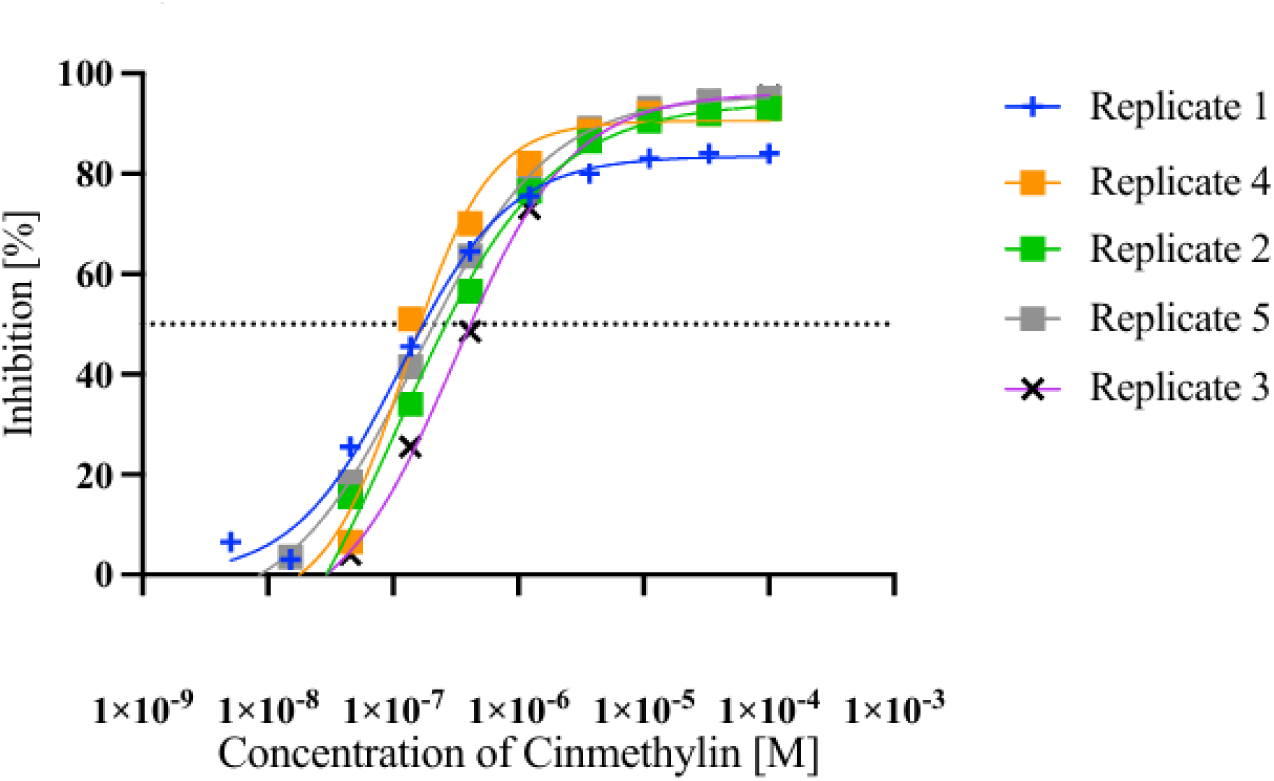
**Determination of IC50 values for *Alopecurus myosuroides* FAT B WT in response to cinmethylin**

### FAT A mutant screening and identification of putative reduced sensitivity candidates

A total of 70 FAT A mutant variants were screened via the acyl-CoA assay. Relative activity in the absence and presence of cinmethylin (10^-4^ M) is shown in (Fig. 7). Residual activity was calculated relative to WT reference. Variants with residual activity <50% of WT were excluded from further sensitivity determination. Among the remaining variants the three mutants R171Q, R171I and R171M showed low inhibition at saturating cinmethylin concentration (24.3%, 15.5%, 11.00% inhibition) while maintaining residual activity >50% of WT (Tab. 3). This indicates a significant reduced sensitivity at saturating inhibitor concentration and highlights residue R171 as a critical position for cinmethylin-mediated inhibition.

**Fig. 7.**
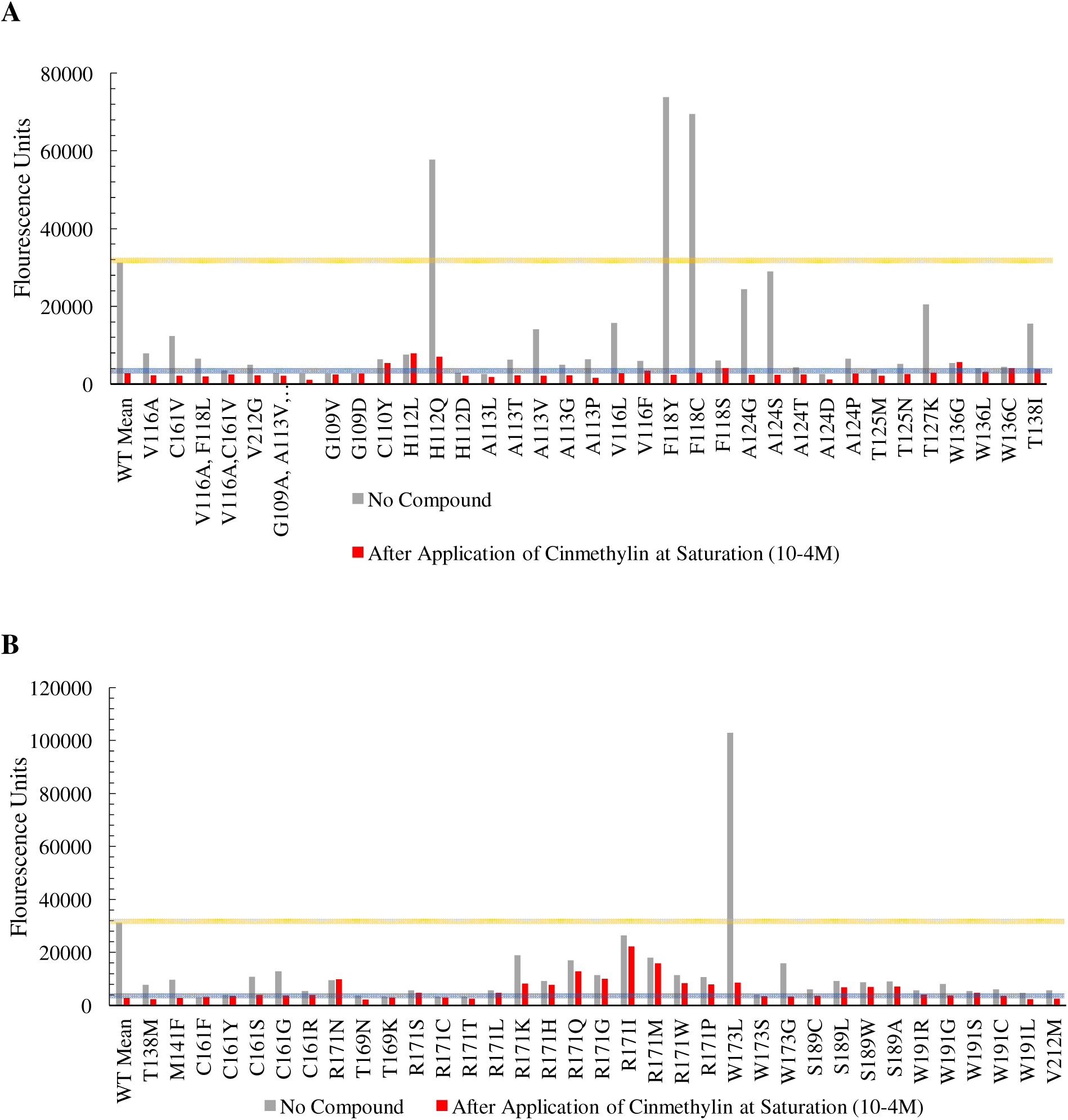
Acyl-CoA Assay of FAT A mutant of *Alopecurus myosuroides* variants before and after cinmethylin treatment. Grey bars represent relative activity without compound application, while red bars show residual activity after application of cinmethylin (10^-4^M). The horizontal lines indicate the reference level corresponding t the WT.

All other variants exceeding the residual activity threshold showed inhibition greater than 50% at 10^-4^ M and were therefore advanced to dose-response analysis for IC_50_ estimation.

### FAT A dose-response analysis and magnitude of sensitivity shifts

Dose-response curves and IC_50_ estimates for prioritized FAT A mutant variants are shown in Fig. 8 and IC_50_ values were compared against WT baseline and summarized as resistance indices (Tab. 3). The largest shift relative to WT was observed for R171K (IC50 1.56 x 10^-5^ M; resistance index 516.6). Additional shifts were observed for W173L (2.05 x 10^-7^ M; resistance index 6.77), H112Q (1.28 x 10^-7^ M; resistance index 4.24) and F118C (8.87 x 10^-8^ M; resistance index 2.94). Approximately twofold increases in IC_50_ were observed for T127K (1.99-fold) and F118Y (2.04-fold). Other variants displayed IC_50_ values comparable to WT. R171Q, R171I and R171M showed the strongest insensitivity levels with no measurable IC50.

**Fig. 8.**
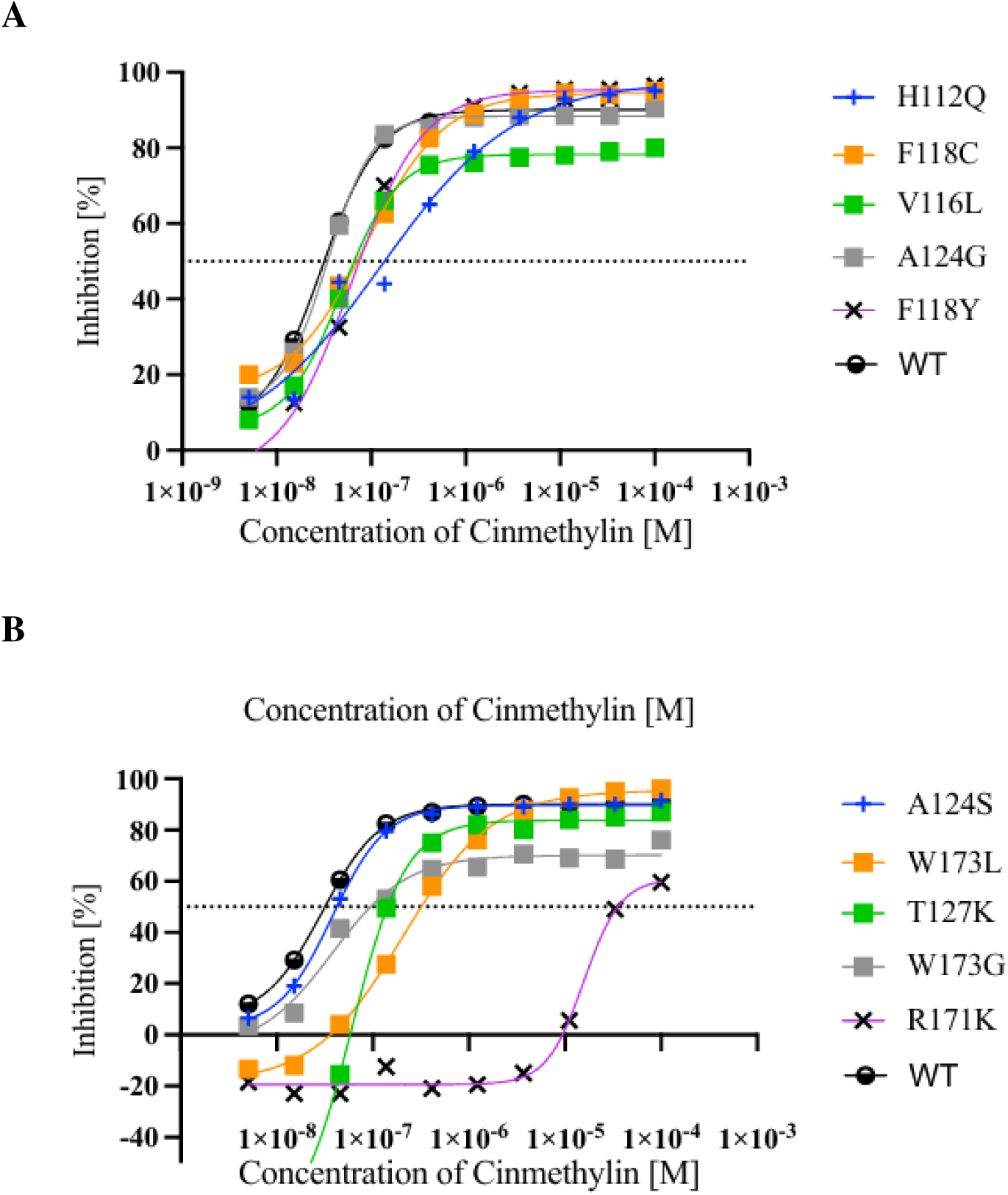
Dose-response curves of *Alopecurus myosuroides* FAT A mutants for (A) H112Q, V116L, F118Y, F118C, A124G and (B) A124S, T127K, W173G, R171K.

These results are consistent with cinmethylin binding to FAT enzymes and suggest that specific substitutions can reduce inhibitor potency, in line with prior structural and biochemical evidence that cinmethylin targets acyl-ACP thioesterases (Campe et al., 2018).

### Codon-based mutation probability assessment for prioritized FAT A variants in A. myosuroides

To contextualize the evolutionary plausibility of the observed sensitivity shifts, codon-level nucleotide polymorphism (NP) requirements were determined for nine prioritized FAT A variants (H112Q, F118Y, F118C, T127K, R171K, R171Q, R171I, R171M, W173L) (Tab. 4). Variants required one, two or three NPs depending on the codon changes. In the defined and simplified model (equal probability of the three alternative bases per position), mutations requiring a single-NP (SNP) represent the most likely class at the population level in this framework, whereas two- and three-NP changes were orders of magnitude less likely to occur. For SNP candidates, codon substitution mappings were generated (Tab. 5). For H112Q, two SNP-derived target codons (CAA or CAG) are possible, leading to an increased within-codon probability (2/9) of reaching a glutamine codon under the model assumptions. While this model does not account for any transition-transversion bias, selection on protein stability etc., it provides a first order ranking of mutational accessibility and supports prioritization of candidates for further monitoring.

**Tab. 4.**
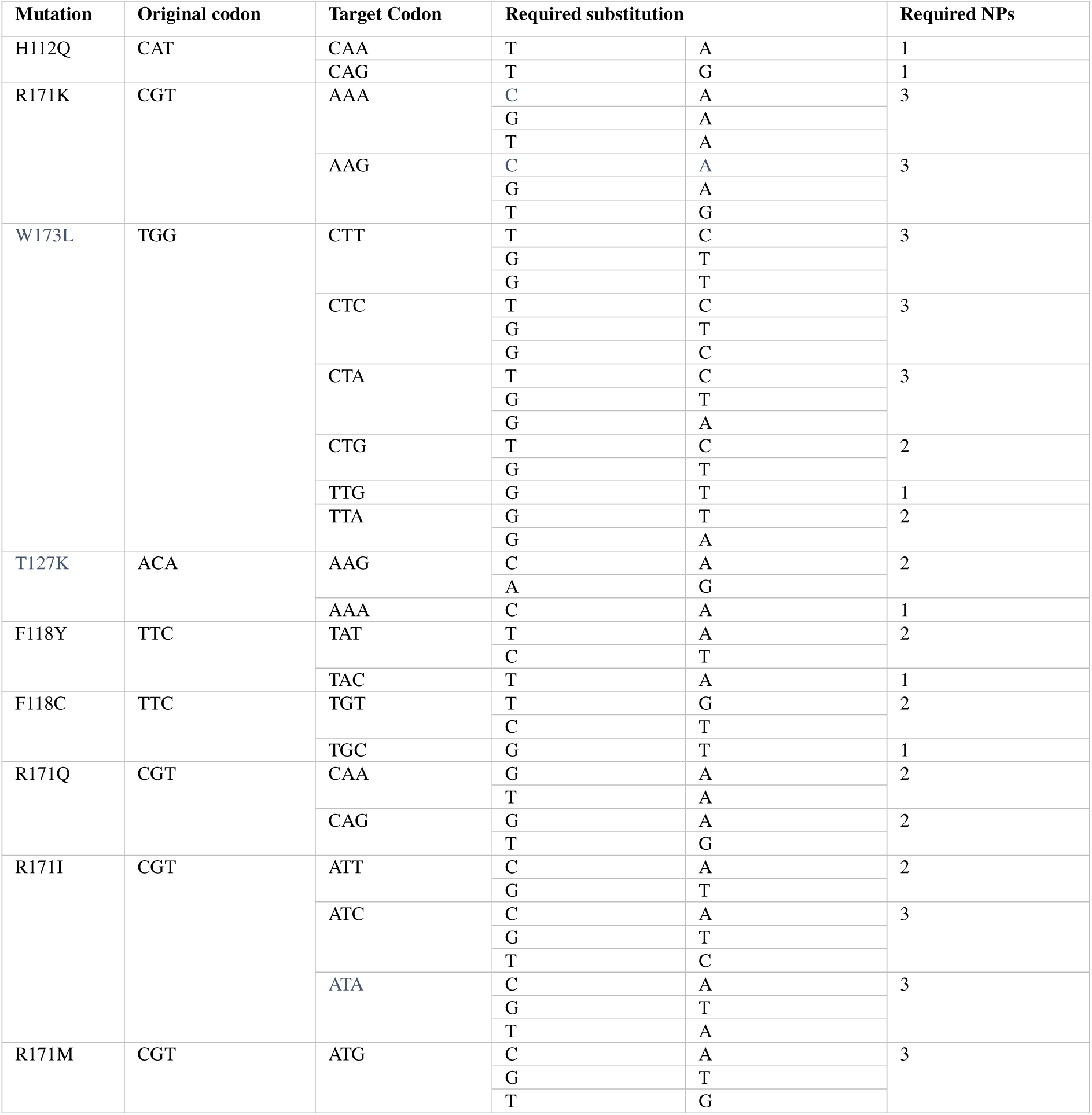
Required nucleotide polymorphisms to reach target codons resulting in respective amino acid exchange in *Alopecurus myosuroides* FAT A.

**Tab. 5.** cDNA-based analysis of *Alopecurus myosuroides* FAT A, showing possible nucleotide substitutions, resulting codons and amino acids and the probability of occurrence of the target mutation

| Mutation | Base | Possible Substitutionen | Resulting Codon | Resulting Aminoacid | Probability [%] |
| --- | --- | --- | --- | --- | --- |
| H112Q | C | A | AAT | N | 22.22 |
|  |  | G | GAT | D |  |
|  |  | T | TAT | Y |  |
|  | A | C | CCT | P |  |
|  |  | G | CGT | R |  |
|  |  | T | CTT | L |  |
|  | T | A | CAA | Q |  |
|  |  | C | CAC | H |  |
|  |  | G | CAG | Q |  |
| F118Y | T | A | ATC | I | 11.11 |
|  |  | C | CTC | L |  |
|  |  | G | GTC | V |  |
|  | T | A | TAC | Y |  |
|  |  | C | TCC | S |  |
|  |  | G | TGC | C |  |
|  | C | A | TTA | L |  |
|  |  | G | TTG | L |  |
|  |  | T | TTT | F |  |
| F118C | T | A | ATC | I | 11.11 |
|  |  | C | CTC | L |  |
|  |  | G | GTC | V |  |
|  | T | A | TAC | Y |  |
|  |  | C | TCC | S |  |
|  |  | G | TGC | C |  |
|  | C | A | TTA | L |  |
|  |  | G | TTG | L |  |
|  |  | T | TTT | F |  |
| A181V | G | A | ACC | T | 11.11 |
|  |  | C | CCC | P |  |
|  |  | T | TCC | S |  |
|  | C | A | GAC | D |  |
|  |  | G | GGC | G |  |
|  |  | T | GTC | V |  |
|  | C | A | GCA | A |  |
|  |  | G | GCG | A |  |
|  |  | T | GCT | A |  |
| V178G | G | A | ATG | M | 11.11 |
|  |  | C | CTG | L |  |
|  |  | T | TTG | L |  |
|  | T | A | GAG | E |  |
|  |  | G | GGG | G |  |
|  |  | C | GCG | A |  |
|  | G | A | GTA | V |  |
|  |  | C | GTC | V |  |
|  |  | T | GTT | V |  |
| P192R | C | A | ACC | T | 11.11 |
|  |  | G | GCC | A |  |
|  |  | T | TCC | S |  |
|  | C | A | CAC | H |  |
|  |  | G | CGC | R |  |
|  |  | T | CTC | L |  |
|  | C | A | CCA | P |  |
|  |  | G | CCG | P |  |
|  |  | T | CCT | P |  |

### Cross-species conservation of FAT A positions in Lolium multiflorum

Because *A. myosuroides* and *L. multiflorum* are agronomically important amongst other things due to their rapid resistance evolution, conservation of prioritized FAT A positions was assessed. FAT A sequences showed high similarity (91.1%). Alignment of the regions of the prioritized positions (H112, R171, W173, T127, F118) indicated conservation of the corresponding amino acids in *L. multiflorum* (Tab. 6). At the cDNA level, codons were identical between both species for H112, R171, W173 and T127, while F118 differed. Consequently, codon-specific NP requirements were recalculated for L. multiflorum for F118Y and F118C and SNP substitution mapping confirmed that model assumptions remained applicable for these targets (Tab. 7). These findings suggest that mutational routes identified in *A. myosuroides* may also be accessible in *L. multiflorum*.

**Tab. 6.** Alignments showing the consensus identity of relevant protein regions in *Alopecurus myosuroides* and *Lolium multiflorum* FAT A.

| Region | Alignment |
| --- | --- |
| H112 | 100<br>110<br>114<br>Val Glu Thr Ile Ala Asn Leu Leu Gln Glu Val Gly Cys Asn His A<br>Consensus Identity<br>Val Glu Thr Ile Ala Asn Leu Leu Gln Glu Val Gly Cys Asn His A<br>Val Glu Thr Ile Ala Asn Leu Leu Gln Glu Val Gly Cys Asn His A<br>1. ALOMY F...<br>2. Lolium F...<br>112 aa & 2 gaps<sup>x</sup> |
| R171 | 160<br>170<br>173<br>Ile Glu Thr Trp Cys Gln Ala Asp Gly Arg Ile Gly Thr Arg Arg A<br>Consensus Identity<br>Ile Glu Thr Trp Cys Gln Ala Asp Gly Arg Ile Gly Thr Arg Arg A<br>Ile Glu Thr Trp Cys Gln Ala Asp Gly Arg Ile Gly Thr Arg Arg A<br>1. ALOMY F...<br>2. Lolium F...<br>171 aa & 2 gaps<sup>x</sup> |
| W173 | 160<br>170<br>175<br>Glu Thr Trp Cys Gln Ala Asp Gly Arg Ile Gly Thr Arg Arg Asp Trp<br>Consensus Identity<br>Glu Thr Trp Cys Gln Ala Asp Gly Arg Ile Gly Thr Arg Arg Asp Trp<br>Glu Thr Trp Cys Gln Ala Asp Gly Arg Ile Gly Thr Arg Arg Asp Trp<br>1. ALOMY F...<br>2. Lolium F...<br>173 aa & 2 gaps<sup>x</sup> |
| T127 | 120<br>129<br>Asn His Ala Gln Ser Val Gly Phe Ser Thr Asp Gly Phe Ala Thr Thr Thr<br>Consensus Identity<br>Asn His Ala Gln Ser Val Gly Phe Ser Thr Asp Gly Phe Ala Thr Thr Thr<br>Asn His Ala Gln Ser Val Gly Phe Ser Thr Asp Gly Phe Ala Thr Thr Thr<br>1. ALOMY F...<br>2. Lolium F...<br>127 aa & 2 gaps<sup>x</sup> |
| F118 | 110<br>120<br>Ala Asn Leu Leu Gln Glu Val Gly Cys Asn His Ala Gln Ser Val Gly Phe S<br>Consensus Identity<br>Ala Asn Leu Leu Gln Glu Val Gly Cys Asn His Ala Gln Ser Val Gly Phe S<br>Ala Asn Leu Leu Gln Glu Val Gly Cys Asn His Ala Gln Ser Val Gly Phe S<br>1. ALOMY F...<br>2. Lolium F...<br>118 aa & 2 gaps<sup>x</sup> |

**Tab. 7.** cDNA-based analysis of *Lolium multiflorum* FAT A and B, showing possible nucleotide substitutions, resulting codons and amino acids and the probability of occurrence of the target mutation

| Mutation | Base | Possible Substitutionen | Resulting Codon | Resulting Aminoacid | Probability [%] |
| --- | --- | --- | --- | --- | --- |
| FAT A |  |  |  |  |  |
| F118Y | T | A | ATT | I | 11,11 |
|  |  | C | CTT | L |  |
|  |  | G | GTT | V |  |
|  | T | A | TAT | Y |  |
|  |  | C | TCT | S |  |
|  |  | G | TGT | C |  |
|  | T | A | TTA | L |  |
|  |  | C | TTC | F |  |
|  |  | G | TTG | L |  |
| F118Y | T | A | ATT | I | 11,11 |
|  |  | C | CTT | L |  |
|  |  | G | GTT | V |  |
|  | T | A | TAT | Y |  |
|  |  | C | TCT | S |  |
|  |  | G | TGT | C |  |
|  | T | A | TTA | L |  |
|  |  | C | TTC | F |  |
|  |  | G | TTG | L |  |
| FAT B |  |  |  |  |  |
| A181V | G | A | ACT | T | 11,11 |
|  |  | C | CCT | P |  |
|  |  | T | TCT | S |  |
|  | C | A | GAT | D |  |
|  |  | G | GGT | G |  |
|  |  | T | GTT | V |  |
|  | T | A | GCA | A |  |
|  |  | G | GCG | A |  |
|  |  | C | GCC | A |  |
| P192R | C | A | ACG | T | 11,11 |
|  |  | G | GCG | A |  |
|  |  | T | TCG | S |  |
|  | C | A | CAG | Q |  |
|  |  | G | CGG | R |  |
|  |  | T | CTG | L |  |
|  | G | A | CCA | P |  |
|  |  | C | CCC | P |  |
|  |  | T | CCT | P |  |

### FAT B mutant screening and dose response analysis

A total of 34 FAT B mutant variants were screened via the acyl-CoA assay (Fig. 9). Variants with residual activity <50% of WT were excluded from further sensitivity classification. The remaining mutants and their inhibition values are summarized in Tab. 3. All listed variants exceeded 50% inhibition at 10^-4^ M were advanced to dose-response analysis (Fig. 10). IC_50_ values are shown in Tab. 3 as well as resistance indices relative to WT. The largest shift was observed for P192R (IC_50_ 1.60 x 10^-6^ M; resistance index: 9.76). V178G (IC_50_ 6.24 x 10^-7^ M; resistance index: 3.80) and A190E (IC_50_ 4.15 x 10^-7^ M; resistance index: 2.53) also showed increased IC_50_ values relative to WT. A181V displayed an approximately twofold increase (1.93-fold). In contrast, V226C and the double mutant A181V&V226C exhibited lower IC_50_ estimates than WT, which indicates that not all substitutions reduce cinmethylin sensitivity and that certain changes may increase apparent susceptibility under the assay conditions.

**Fig. 9.**
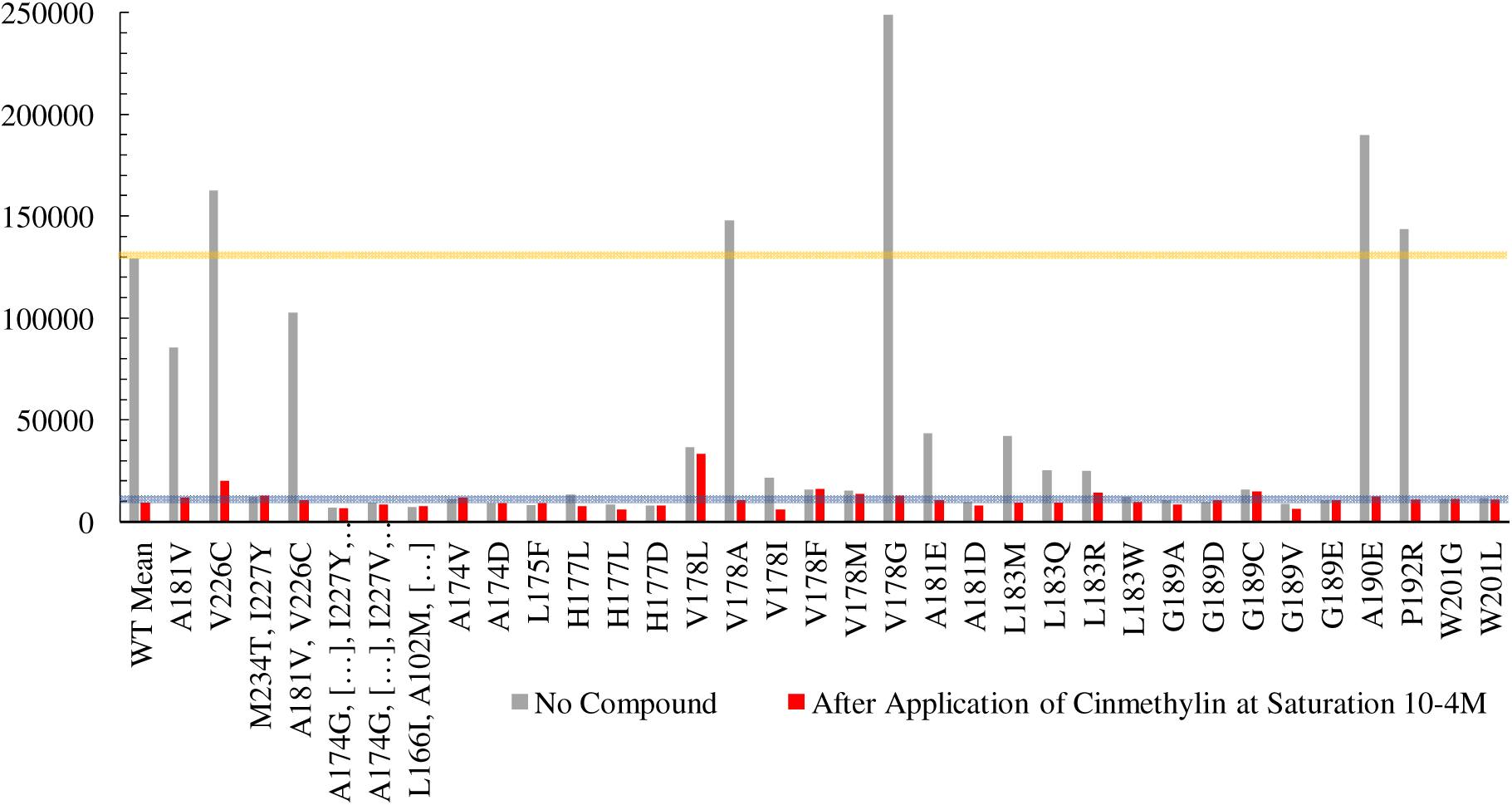
Acyl-CoA Assay of *Alopecurus myosuroides* FAT B mutant variants before and after cinmethylin application. Grey bars represent relative activity without compound application, while red bars show activity after application of cinmethylin (10^-4^M). The horizontal lines indicate the reference level corresponding to the WT.

**Fig. 10.**
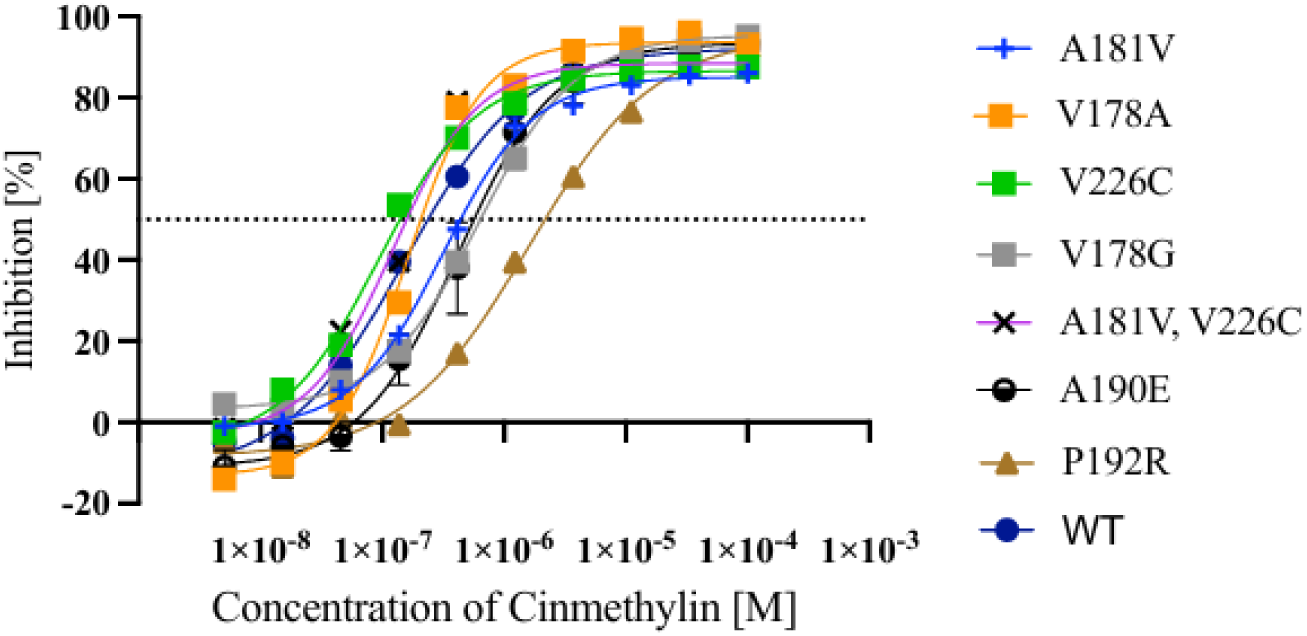
**Dose-response curves of *Alopecurus myosuroides* FAT B mutants.**

### Codon-based mutation probability assessment for prioritized FAT B variants and cross-species context

For the prioritized FAT B variants (A181V, V178G, A190E, P192R), required NPs to reach target codons *in A. myosuroides* were determined (Tab. 8). SNP substitution profiles were generated for selected variants (Tab. 9) yielding a target-codon probability of 1/9 (11.11%) for the analyzed variants under model assumptions.

**Tab. 8.** Required nucleotide polymorphisms to reach target codons resulting in respective amino acid exchange in *Alopecurus myosuroides* FAT B.

| Mutation | Original Codon | Target Codon | Required substitution (Before / After) |  | Required NPs |
| --- | --- | --- | --- | --- | --- |
| A181V | GCC | GTG | C | T | 2 |
|  |  |  | C | G |  |
|  |  | GTA | C | T | 2 |
|  |  |  | C | A |  |
|  |  | GTC | C | T | 1 |
|  |  | GTT | C | T | 2 |
|  |  |  | C | T |  |
| V178G | GTG | GGG | T | G | 1 |
|  |  | GGA | T | G | 2 |
|  |  |  | G | A |  |
|  |  | GGC | T | G | 2 |
|  |  |  | G | C |  |
|  |  | GGT | T | G | 2 |
|  |  |  | G | T |  |
| A190E | GCT | GAG | C | A | 2 |
|  |  |  | T | G |  |
|  |  | GAA | C | A | 2 |
|  |  |  | T | A |  |
| P192R | CCC | CGG | C | G | 2 |
|  |  |  | C | G |  |
|  |  | CGA | C | G | 2 |
|  |  |  | C | A |  |
|  |  | CGC | C | G | 1 |
|  |  | CGT | C | G | 2 |
|  |  |  | C | T |  |
|  |  | AGG | C | A | 3 |
|  |  |  | C | G |  |
|  |  |  | C | G |  |
|  |  | AGA | C | A | 3 |
|  |  |  | C | G |  |
|  |  |  | C | A |  |

**Tab. 9.** cDNA-based analysis of *Alopecurus myosuroides* FAT A, showing possible nucleotide substitutions, resulting codons and amino acids and the probability of occurrence of the target mutation

| Mutation | Base | Possible Substitutionen | Resulting Codon | Resulting Aminoacid | Probability [%] |
| --- | --- | --- | --- | --- | --- |
| A181V | G | A | ACC | T | 11,11 |
|  |  | C | CCC | P |  |
|  |  | T | TCC | S |  |
|  | C | A | GAC | D |  |
|  |  | G | GGC | G |  |
|  |  | T | GTC | V |  |
|  | C | A | GCA | A |  |
|  |  | G | GCG | A |  |
|  |  | T | GCT | A |  |
| V178G | G | A | ATG | M | 11,11 |
|  |  | C | CTG | L |  |
|  |  | T | TTG | L |  |
|  | T | A | GAG | E |  |
|  |  | G | GGG | G |  |
|  |  | C | GCG | A |  |
|  | G | A | GTA | V |  |
|  |  | C | GTC | V |  |
|  |  | T | GTT | V |  |
| P192R | C | A | ACC | T | 11,11 |
|  |  | G | GCC | A |  |
|  |  | T | TCC | S |  |
|  | C | A | CAC | H |  |
|  |  | G | CGC | R |  |
|  |  | T | CTC | L |  |
|  | C | A | CCA | P |  |
|  |  | G | CCG | P |  |
|  |  | T | CCT | P |  |

*A. myosuroides* and *L. multiflorum* were highly similar for FAT B (93.2%). Alignment indicated conservation of the critical amino acids (A181, V178, A190, P192) across species (Tab. 10) Codon comparisons revealed differences for A181 and P192 which leads to recalculation of NP requirements (Tab. 7).

**Tab. 10.** Alignments showing the consensus identity of relevant protein regions in *Alopecurus myosuroides* and *Lolium multiflorum* FAT.

| Region | Alignment |
| --- | --- |
| A181 | Consensus Identity<br>170 180 182<br>et Asn His Met Gln Glu Thr Ala Leu Asn His Val Lys Thr Ala<br>1. ALOMY F... et Asn His Met Gln Glu Thr Ala Leu Asn His Val Lys Thr Ala<br>2. Lolium F... et Asn His Met Gln Glu Thr Ala Leu Asn His Val Lys Thr Ala<br>181 aa & 1 gap<sup>x</sup> |
| V178 | Consensus Identity<br>170 179<br>Thr Leu Met Asn His Met Gln Glu Thr Ala Leu Asn His Val<br>1. ALOMY F... Thr Leu Met Asn His Met Gln Glu Thr Ala Leu Asn His Val<br>2. Lolium F... Thr Leu Met Asn His Met Gln Glu Thr Ala Leu Asn His Val<br>178 aa & 1 gap<sup>x</sup> |
| A190 | Consensus Identity<br>180 191<br>Leu Asn His Val Lys Thr Ala Gly Leu Leu Gly Asp Gly Phe Gly Ala T<br>1. ALOMY F... Leu Asn His Val Lys Thr Ala Gly Leu Leu Gly Asp Gly Phe Gly Ala T<br>2. Lolium F... Leu Asn His Val Lys Thr Ala Gly Leu Leu Gly Asp Gly Phe Gly Ala T<br>190 aa & 1 gap<sup>x</sup> |
| P192 | Consensus Identity<br>180 190 193<br>s Val Lys Thr Ala Gly Leu Leu Gly Asp Gly Phe Gly Ala Thr Pro<br>1. ALOMY F... s Val Lys Thr Ala Gly Leu Leu Gly Asp Gly Phe Gly Ala Thr Pro<br>2. Lolium F... s Val Lys Thr Ala Gly Leu Leu Gly Asp Gly Phe Gly Ala Thr Pro<br>192 aa & 1 gap<sup>x</sup> |

### Implications for cinmethylin durability and resistance risk

Herbicide resistance is a major challenge in modern agriculture, particularly in grass weeds with large population sizes and strong selection pressure (Kersten et al., 2023). Cinmethylin represents an important alternative mode of action through inhibition of fatty acid thioesterases, key enzymes in the de novo fatty acid biosynthesis that are essential for plant viability (Campe et al., 2018; Ohlrogge & Browse, 1995).

The findings of this study provide first indications that cinmethylin, despite the absence of reported resistance cases to date, is not immune to potential resistance evolution. Specific amino acid substitutions in the target proteins FAT A and FAT B can reduce cinmethylin sensitivity in vitro, which indicates that target-site resistance is theoretically possible. However, the magnitude of the observed IC_50_ shifts was generally moderate.

In weed species with high population sizes, such as *Alopecurus myosuroides* and *Lolium multiflorum*, the emergence of resistant individuals under repeated cinmethylin selection can not be excluded. The high degree of conservation observed at critical amino acid positions between these species further suggests that resistance-conferring substitutions arising in one species may analogously evolve in another.

At the molecular level, the present screening indicates that specific target-site substitutions in FAT A and B can reduce cinmethylin sensitivity in vitro, with the strongest effects observed at FAT A residue R171 and the FAT B substitution P192R. Structural evidence supports these observations, as cinmethylin binding involves interactions within a defined pocket in which a conserved arginine residue contributes to hydrogen bonding and ligand stabilization (Campe et al., 2018). Substitutions at R171 alter the side-chain geometry and therefore weaken the binding affinity to the compound.

Importantly, among the many mutant variants tested, only a small number retained at least 50% residual enzymatic activity and would therefore be expected to remain viable in plants. While substitutions at R171 exhibited the strongest reduction in cinmethylin binding and therefore represent the greatest theoretical resistance risk, they are extremely unlikely to occur under field conditions, as they would require multiple specific nucleotide polymorphisms. By contrast, the H112Q substitution shows a more realistic change, its effect on cinmethylin binding was comparatively modest.

Overall, this study demonstrates that while cinmethylin represents an important herbicide with a novel mode of action and no reported resistance cases to date, its efficacy could in theory be affected by specific target-site mutations in FAT enzymes, particularly those arising from SNPs. However, the findings clearly demonstrate that while theoretical resistance routes exist, the practical risk remains low, underscoring cinmethylin robustness as a reliable herbicide option. The high sequence conservation between *Alopecurus myosuroides* and *Lolium multiflorum* underscores the agronomic relevance of these findings, as resistance evolution in both species appears possible under selection pressure. This highlights the necessity for proactive, integrated resistance management to safeguard its long-term effectiveness.

### Limitations

Although the results of the study provide important insights, the mutations analyzed were tested only in vitro and their actual impact on resistance in plants remains to be determined. While these provide theoretical insights, they cannot capture the complex interplay of metabolism, fitness costs and selection pressure occurring in field populations. Future work should therefore include ectopic expression in model species such as *Arabidopsis thaliana* to assess in vivo functionality.

The codon-based probability model applied in this study represents a strongly simplified framework. It assumes equal mutation probabilities for all base substitutions and does not account for biological realities. As a result, the calculated probabilities must be interpreted as approximations rather than absolute values.

Nevertheless, the combined analysis presented here provides a valuable framework for assessing the potential risk of resistance evolution against cinmethylin.

## Acknowledgments

The author thanks Dr. Aimone Porri (BASF Agricultural Center, Limburgerhof, Germany) for supervision and guidance throughout the research project and department of phytopathology at RPTU Kaiserslautern for academic support. Technical assistance from BASF laboratory staff is gratefully acknowledged.

## Supplementary data

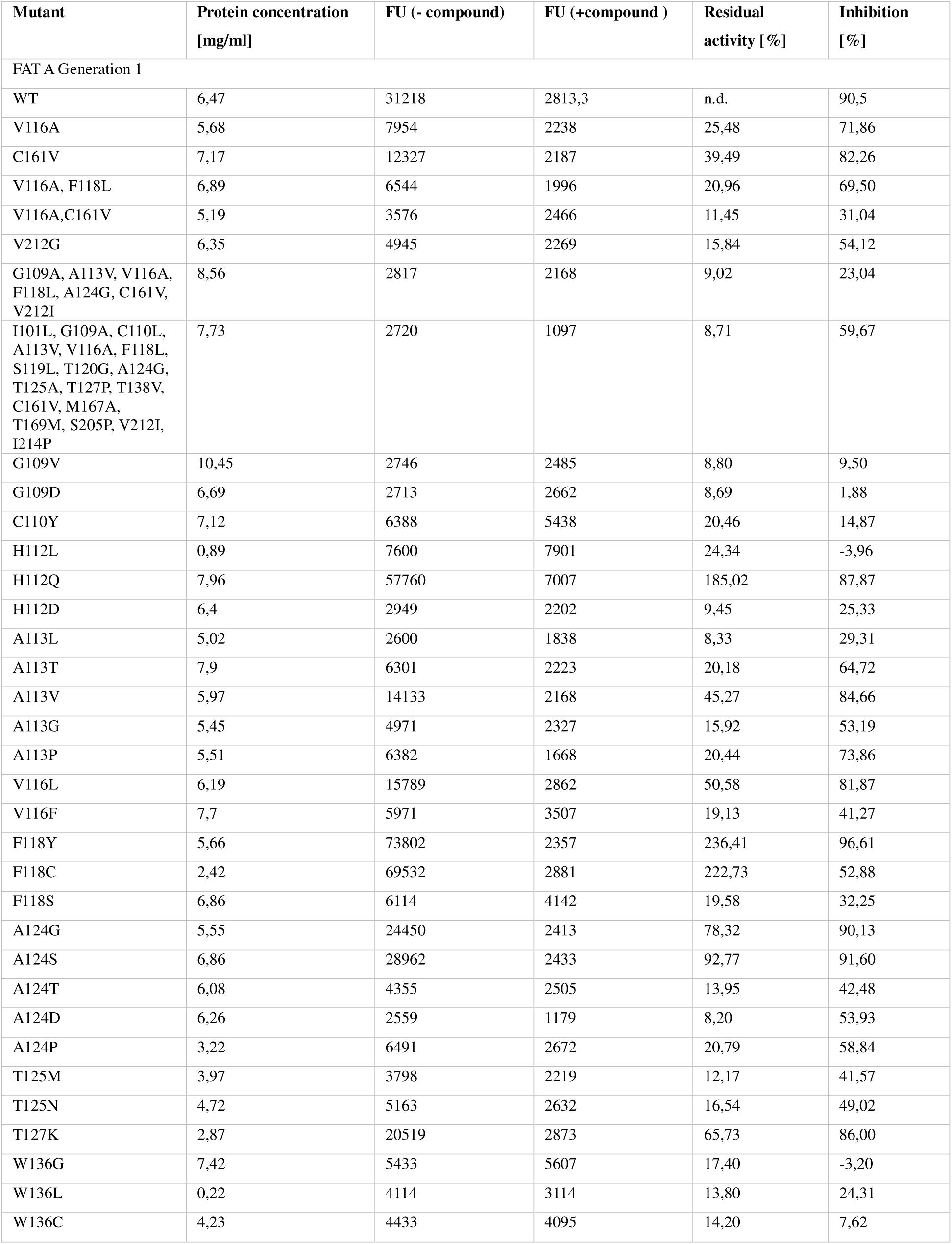
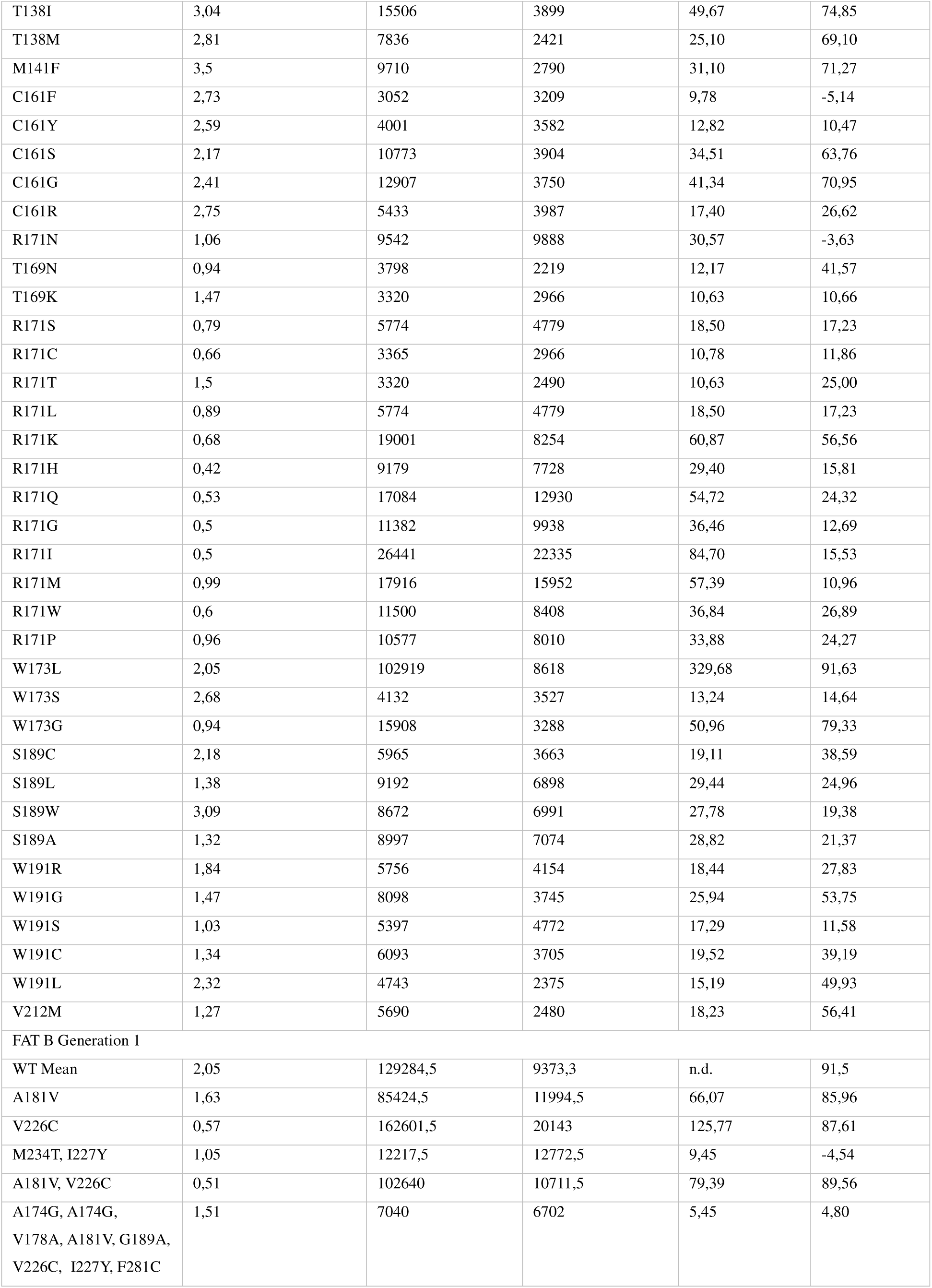
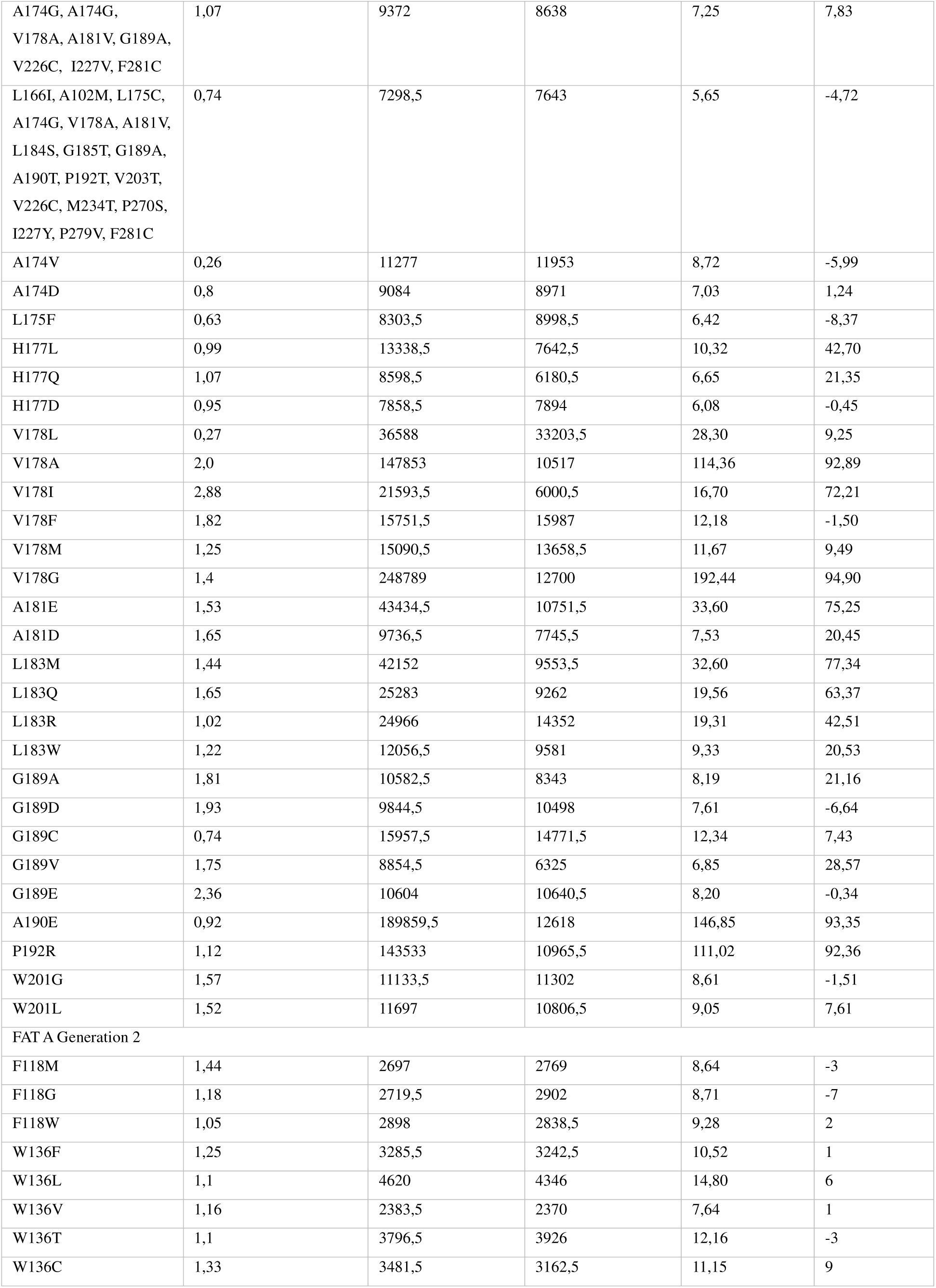
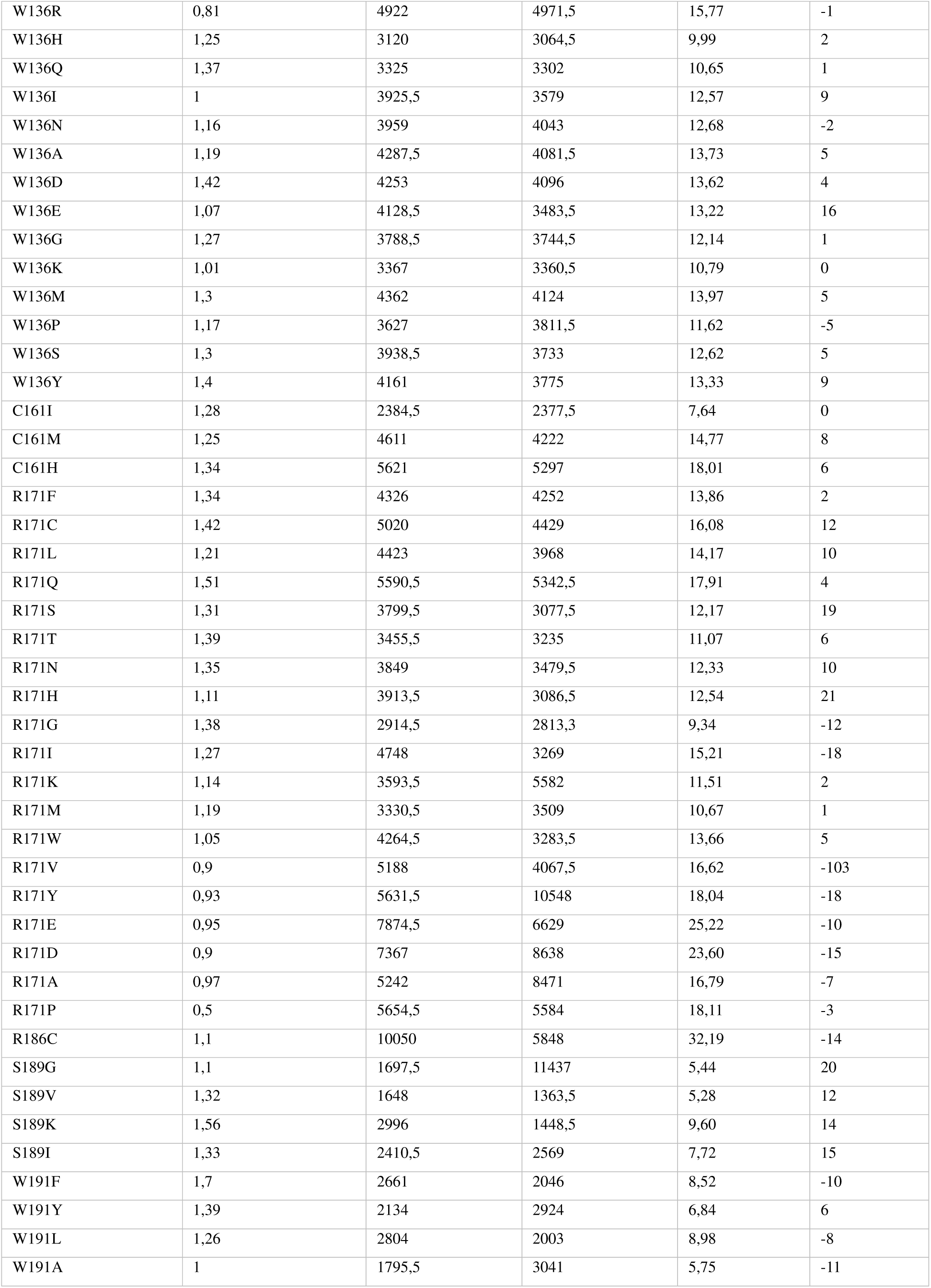
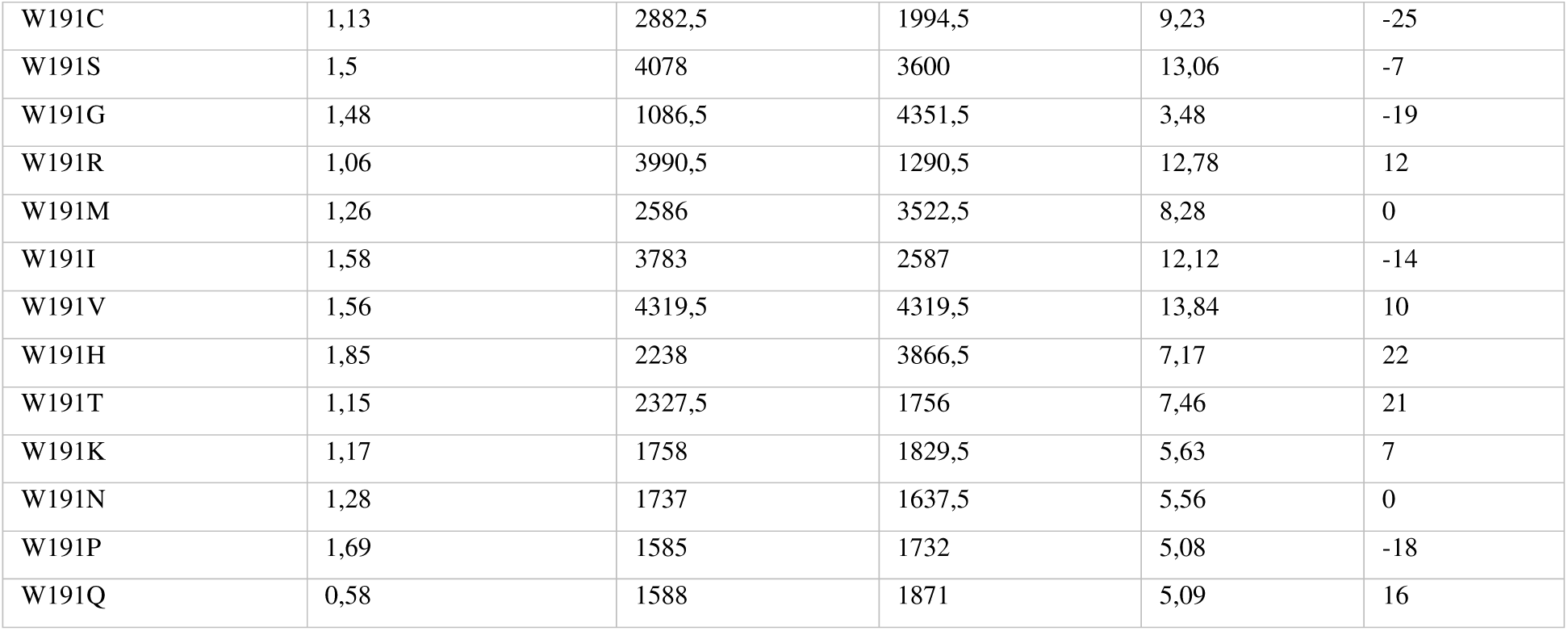

| Mutant | Protein concentration<br>[mg/ml] | FU (- compound) | FU (+compound ) | Residual<br>activity [%] | Inhibition<br>[%] |
| --- | --- | --- | --- | --- | --- |
| FAT A Generation 1 |  |  |  |  |  |
| WT | 6,47 | 31218 | 2813,3 | n.d. | 90,5 |
| V116A | 5,68 | 7954 | 2238 | 25,48 | 71,86 |
| C161V | 7,17 | 12327 | 2187 | 39,49 | 82,26 |
| V116A, F118L | 6,89 | 6544 | 1996 | 20,96 | 69,50 |
| V116A,C161V | 5,19 | 3576 | 2466 | 11,45 | 31,04 |
| V212G | 6,35 | 4945 | 2269 | 15,84 | 54,12 |
| G109A, A113V, V116A,<br>F118L, A124G, C161V,<br>V212I | 8,56 | 2817 | 2168 | 9,02 | 23,04 |
| I101L, G109A, C110L,<br>A113V, V116A, F118L,<br>S119L, T120G, A124G,<br>T125A, T127P, T138V,<br>C161V, M167A,<br>T169M, S205P, V212I,<br>I214P | 7,73 | 2720 | 1097 | 8,71 | 59,67 |
| G109V | 10,45 | 2746 | 2485 | 8,80 | 9,50 |
| G109D | 6,69 | 2713 | 2662 | 8,69 | 1,88 |
| C110Y | 7,12 | 6388 | 5438 | 20,46 | 14,87 |
| H112L | 0,89 | 7600 | 7901 | 24,34 | -3,96 |
| H112Q | 7,96 | 57760 | 7007 | 185,02 | 87,87 |
| H112D | 6,4 | 2949 | 2202 | 9,45 | 25,33 |
| A113L | 5,02 | 2600 | 1838 | 8,33 | 29,31 |
| A113T | 7,9 | 6301 | 2223 | 20,18 | 64,72 |
| A113V | 5,97 | 14133 | 2168 | 45,27 | 84,66 |
| A113G | 5,45 | 4971 | 2327 | 15,92 | 53,19 |
| A113P | 5,51 | 6382 | 1668 | 20,44 | 73,86 |
| V116L | 6,19 | 15789 | 2862 | 50,58 | 81,87 |
| V116F | 7,7 | 5971 | 3507 | 19,13 | 41,27 |
| F118Y | 5,66 | 73802 | 2357 | 236,41 | 96,61 |
| F118C | 2,42 | 69532 | 2881 | 222,73 | 52,88 |
| F118S | 6,86 | 6114 | 4142 | 19,58 | 32,25 |
| A124G | 5,55 | 24450 | 2413 | 78,32 | 90,13 |
| A124S | 6,86 | 28962 | 2433 | 92,77 | 91,60 |
| A124T | 6,08 | 4355 | 2505 | 13,95 | 42,48 |
| A124D | 6,26 | 2559 | 1179 | 8,20 | 53,93 |
| A124P | 3,22 | 6491 | 2672 | 20,79 | 58,84 |
| T125M | 3,97 | 3798 | 2219 | 12,17 | 41,57 |
| T125N | 4,72 | 5163 | 2632 | 16,54 | 49,02 |
| T127K | 2,87 | 20519 | 2873 | 65,73 | 86,00 |
| W136G | 7,42 | 5433 | 5607 | 17,40 | -3,20 |
| W136L | 0,22 | 4114 | 3114 | 13,80 | 24,31 |
| W136C | 4,23 | 4433 | 4095 | 14,20 | 7,62 |
| T138I | 3,04 | 15506 | 3899 | 49,67 | 74,85 |
| T138M | 2,81 | 7836 | 2421 | 25,10 | 69,10 |
| M141F | 3,5 | 9710 | 2790 | 31,10 | 71,27 |
| C161F | 2,73 | 3052 | 3209 | 9,78 | -5,14 |
| C161Y | 2,59 | 4001 | 3582 | 12,82 | 10,47 |
| C161S | 2,17 | 10773 | 3904 | 34,51 | 63,76 |
| C161G | 2,41 | 12907 | 3750 | 41,34 | 70,95 |
| C161R | 2,75 | 5433 | 3987 | 17,40 | 26,62 |
| R171N | 1,06 | 9542 | 9888 | 30,57 | -3,63 |
| T169N | 0,94 | 3798 | 2219 | 12,17 | 41,57 |
| T169K | 1,47 | 3320 | 2966 | 10,63 | 10,66 |
| R171S | 0,79 | 5774 | 4779 | 18,50 | 17,23 |
| R171C | 0,66 | 3365 | 2966 | 10,78 | 11,86 |
| R171T | 1,5 | 3320 | 2490 | 10,63 | 25,00 |
| R171L | 0,89 | 5774 | 4779 | 18,50 | 17,23 |
| R171K | 0,68 | 19001 | 8254 | 60,87 | 56,56 |
| R171H | 0,42 | 9179 | 7728 | 29,40 | 15,81 |
| R171Q | 0,53 | 17084 | 12930 | 54,72 | 24,32 |
| R171G | 0,5 | 11382 | 9938 | 36,46 | 12,69 |
| R171I | 0,5 | 26441 | 22335 | 84,70 | 15,53 |
| R171M | 0,99 | 17916 | 15952 | 57,39 | 10,96 |
| R171W | 0,6 | 11500 | 8408 | 36,84 | 26,89 |
| R171P | 0,96 | 10577 | 8010 | 33,88 | 24,27 |
| W173L | 2,05 | 102919 | 8618 | 329,68 | 91,63 |
| W173S | 2,68 | 4132 | 3527 | 13,24 | 14,64 |
| W173G | 0,94 | 15908 | 3288 | 50,96 | 79,33 |
| S189C | 2,18 | 5965 | 3663 | 19,11 | 38,59 |
| S189L | 1,38 | 9192 | 6898 | 29,44 | 24,96 |
| S189W | 3,09 | 8672 | 6991 | 27,78 | 19,38 |
| S189A | 1,32 | 8997 | 7074 | 28,82 | 21,37 |
| W191R | 1,84 | 5756 | 4154 | 18,44 | 27,83 |
| W191G | 1,47 | 8098 | 3745 | 25,94 | 53,75 |
| W191S | 1,03 | 5397 | 4772 | 17,29 | 11,58 |
| W191C | 1,34 | 6093 | 3705 | 19,52 | 39,19 |
| W191L | 2,32 | 4743 | 2375 | 15,19 | 49,93 |
| V212M | 1,27 | 5690 | 2480 | 18,23 | 56,41 |
| FAT B Generation 1 |  |  |  |  |  |
| WT Mean | 2,05 | 129284,5 | 9373,3 | n.d. | 91,5 |
| A181V | 1,63 | 85424,5 | 11994,5 | 66,07 | 85,96 |
| V226C | 0,57 | 162601,5 | 20143 | 125,77 | 87,61 |
| M234T, I227Y | 1,05 | 12217,5 | 12772,5 | 9,45 | -4,54 |
| A181V, V226C | 0,51 | 102640 | 10711,5 | 79,39 | 89,56 |
| A174G, A174G,<br>V178A, A181V, G189A,<br>V226C, I227Y, F281C | 1,51 | 7040 | 6702 | 5,45 | 4,80 |
| A174G, A174G,<br>V178A, A181V, G189A,<br>V226C, I227V, F281C | 1,07 | 9372 | 8638 | 7,25 | 7,83 |
| L166I, A102M, L175C,<br>A174G, V178A, A181V,<br>L184S, G185T, G189A,<br>A190T, P192T, V203T,<br>V226C, M234T, P270S,<br>I227Y, P279V, F281C | 0,74 | 7298,5 | 7643 | 5,65 | -4,72 |
| A174V | 0,26 | 11277 | 11953 | 8,72 | -5,99 |
| A174D | 0,8 | 9084 | 8971 | 7,03 | 1,24 |
| L175F | 0,63 | 8303,5 | 8998,5 | 6,42 | -8,37 |
| H177L | 0,99 | 13338,5 | 7642,5 | 10,32 | 42,70 |
| H177Q | 1,07 | 8598,5 | 6180,5 | 6,65 | 21,35 |
| H177D | 0,95 | 7858,5 | 7894 | 6,08 | -0,45 |
| V178L | 0,27 | 36588 | 33203,5 | 28,30 | 9,25 |
| V178A | 2,0 | 147853 | 10517 | 114,36 | 92,89 |
| V178I | 2,88 | 21593,5 | 6000,5 | 16,70 | 72,21 |
| V178F | 1,82 | 15751,5 | 15987 | 12,18 | -1,50 |
| V178M | 1,25 | 15090,5 | 13658,5 | 11,67 | 9,49 |
| V178G | 1,4 | 248789 | 12700 | 192,44 | 94,90 |
| A181E | 1,53 | 43434,5 | 10751,5 | 33,60 | 75,25 |
| A181D | 1,65 | 9736,5 | 7745,5 | 7,53 | 20,45 |
| L183M | 1,44 | 42152 | 9553,5 | 32,60 | 77,34 |
| L183Q | 1,65 | 25283 | 9262 | 19,56 | 63,37 |
| L183R | 1,02 | 24966 | 14352 | 19,31 | 42,51 |
| L183W | 1,22 | 12056,5 | 9581 | 9,33 | 20,53 |
| G189A | 1,81 | 10582,5 | 8343 | 8,19 | 21,16 |
| G189D | 1,93 | 9844,5 | 10498 | 7,61 | -6,64 |
| G189C | 0,74 | 15957,5 | 14771,5 | 12,34 | 7,43 |
| G189V | 1,75 | 8854,5 | 6325 | 6,85 | 28,57 |
| G189E | 2,36 | 10604 | 10640,5 | 8,20 | -0,34 |
| A190E | 0,92 | 189859,5 | 12618 | 146,85 | 93,35 |
| P192R | 1,12 | 143533 | 10965,5 | 111,02 | 92,36 |
| W201G | 1,57 | 11133,5 | 11302 | 8,61 | -1,51 |
| W201L | 1,52 | 11697 | 10806,5 | 9,05 | 7,61 |
| FAT A Generation 2 |  |  |  |  |  |
| F118M | 1,44 | 2697 | 2769 | 8,64 | -3 |
| F118G | 1,18 | 2719,5 | 2902 | 8,71 | -7 |
| F118W | 1,05 | 2898 | 2838,5 | 9,28 | 2 |
| W136F | 1,25 | 3285,5 | 3242,5 | 10,52 | 1 |
| W136L | 1,1 | 4620 | 4346 | 14,80 | 6 |
| W136V | 1,16 | 2383,5 | 2370 | 7,64 | 1 |
| W136T | 1,1 | 3796,5 | 3926 | 12,16 | -3 |
| W136C | 1,33 | 3481,5 | 3162,5 | 11,15 | 9 |
| W136R | 0,81 | 4922 | 4971,5 | 15,77 | -1 |
| W136H | 1,25 | 3120 | 3064,5 | 9,99 | 2 |
| W136Q | 1,37 | 3325 | 3302 | 10,65 | 1 |
| W136I | 1 | 3925,5 | 3579 | 12,57 | 9 |
| W136N | 1,16 | 3959 | 4043 | 12,68 | -2 |
| W136A | 1,19 | 4287,5 | 4081,5 | 13,73 | 5 |
| W136D | 1,42 | 4253 | 4096 | 13,62 | 4 |
| W136E | 1,07 | 4128,5 | 3483,5 | 13,22 | 16 |
| W136G | 1,27 | 3788,5 | 3744,5 | 12,14 | 1 |
| W136K | 1,01 | 3367 | 3360,5 | 10,79 | 0 |
| W136M | 1,3 | 4362 | 4124 | 13,97 | 5 |
| W136P | 1,17 | 3627 | 3811,5 | 11,62 | -5 |
| W136S | 1,3 | 3938,5 | 3733 | 12,62 | 5 |
| W136Y | 1,4 | 4161 | 3775 | 13,33 | 9 |
| C161I | 1,28 | 2384,5 | 2377,5 | 7,64 | 0 |
| C161M | 1,25 | 4611 | 4222 | 14,77 | 8 |
| C161H | 1,34 | 5621 | 5297 | 18,01 | 6 |
| R171F | 1,34 | 4326 | 4252 | 13,86 | 2 |
| R171C | 1,42 | 5020 | 4429 | 16,08 | 12 |
| R171L | 1,21 | 4423 | 3968 | 14,17 | 10 |
| R171Q | 1,51 | 5590,5 | 5342,5 | 17,91 | 4 |
| R171S | 1,31 | 3799,5 | 3077,5 | 12,17 | 19 |
| R171T | 1,39 | 3455,5 | 3235 | 11,07 | 6 |
| R171N | 1,35 | 3849 | 3479,5 | 12,33 | 10 |
| R171H | 1,11 | 3913,5 | 3086,5 | 12,54 | 21 |
| R171G | 1,38 | 2914,5 | 2813,3 | 9,34 | -12 |
| R171I | 1,27 | 4748 | 3269 | 15,21 | -18 |
| R171K | 1,14 | 3593,5 | 5582 | 11,51 | 2 |
| R171M | 1,19 | 3330,5 | 3509 | 10,67 | 1 |
| R171W | 1,05 | 4264,5 | 3283,5 | 13,66 | 5 |
| R171V | 0,9 | 5188 | 4067,5 | 16,62 | -103 |
| R171Y | 0,93 | 5631,5 | 10548 | 18,04 | -18 |
| R171E | 0,95 | 7874,5 | 6629 | 25,22 | -10 |
| R171D | 0,9 | 7367 | 8638 | 23,60 | -15 |
| R171A | 0,97 | 5242 | 8471 | 16,79 | -7 |
| R171P | 0,5 | 5654,5 | 5584 | 18,11 | -3 |
| R186C | 1,1 | 10050 | 5848 | 32,19 | -14 |
| S189G | 1,1 | 1697,5 | 11437 | 5,44 | 20 |
| S189V | 1,32 | 1648 | 1363,5 | 5,28 | 12 |
| S189K | 1,56 | 2996 | 1448,5 | 9,60 | 14 |
| S189I | 1,33 | 2410,5 | 2569 | 7,72 | 15 |
| W191F | 1,7 | 2661 | 2046 | 8,52 | -10 |
| W191Y | 1,39 | 2134 | 2924 | 6,84 | 6 |
| W191L | 1,26 | 2804 | 2003 | 8,98 | -8 |
| W191A | 1 | 1795,5 | 3041 | 5,75 | -11 |
| W191C | 1,13 | 2882,5 | 1994,5 | 9,23 | -25 |
| W191S | 1,5 | 4078 | 3600 | 13,06 | -7 |
| W191G | 1,48 | 1086,5 | 4351,5 | 3,48 | -19 |
| W191R | 1,06 | 3990,5 | 1290,5 | 12,78 | 12 |
| W191M | 1,26 | 2586 | 3522,5 | 8,28 | 0 |
| W191I | 1,58 | 3783 | 2587 | 12,12 | -14 |
| W191V | 1,56 | 4319,5 | 4319,5 | 13,84 | 10 |
| W191H | 1,85 | 2238 | 3866,5 | 7,17 | 22 |
| W191T | 1,15 | 2327,5 | 1756 | 7,46 | 21 |
| W191K | 1,17 | 1758 | 1829,5 | 5,63 | 7 |
| W191N | 1,28 | 1737 | 1637,5 | 5,56 | 0 |
| W191P | 1,69 | 1585 | 1732 | 5,08 | -18 |
| W191Q | 0,58 | 1588 | 1871 | 5,09 | 16 |

